# Proteolytic cleavage and inactivation of the TRMT1 tRNA modification enzyme by SARS-CoV-2 main protease

**DOI:** 10.1101/2023.02.10.527147

**Authors:** Kejia Zhang, Patrick Eldin, Jessica H. Ciesla, Laurence Briant, Jenna M. Lentini, Jillian Ramos, Justin Cobb, Joshua Munger, Dragony Fu

**Author notes:** Corresponding author: Dragony Fu **Email:**.

## Abstract

Nonstructural protein 5 (Nsp5) is the main protease of SARS-CoV-2 that cleaves viral polyproteins into individual polypeptides necessary for viral replication. Here, we show that Nsp5 binds and cleaves human tRNA methyltransferase 1 (TRMT1), a host enzyme required for a prevalent post-transcriptional modification in tRNAs. Human cells infected with SARS-CoV-2 exhibit a decrease in TRMT1 protein levels and TRMT1-catalyzed tRNA modifications, consistent with TRMT1 cleavage and inactivation by Nsp5. Nsp5 cleaves TRMT1 at a specific position that matches the consensus sequence of SARS-CoV-2 polyprotein cleavage sites, and a single mutation within the sequence inhibits Nsp5-dependent proteolysis of TRMT1. The TRMT1 cleavage fragments exhibit altered RNA binding activity and are unable to rescue tRNA modification in TRMT1-deficient human cells. Compared to wildtype human cells, TRMT1-deficient human cells infected with SARS-CoV-2 exhibit reduced levels of intracellular viral RNA. These findings provide evidence that Nsp5-dependent cleavage of TRMT1 and perturbation of tRNA modification patterns contribute to the cellular pathogenesis of SARS-CoV-2 infection.

## Introduction

Severe acute respiratory syndrome coronavirus 2 (SARS-CoV-2) is an enveloped, single-stranded RNA virus that is the causative agent of the COVID-19 pandemic (Lamers & Haagmans, 2022; Merad *et al*, 2022). SARS-CoV-2 is a member of the Betacoronavirus genus of the Coronaviridae family, which includes the severe acute respiratory syndrome coronavirus 1 (SARS-CoV-1) and Middle East respiratory syndrome coronavirus (MERS-CoV) (reviewed in (Hartenian *et al*, 2020)). SARS-CoV-2 primarily targets the human respiratory tract and lungs, which can clinically manifest as an acute respiratory distress syndrome. Once delivered into the host cell, the incoming positive-strand viral RNA genome is first translated by host ribosomes into two overlapping polyproteins, pp1a and pp1ab. Viral polyproteins are then proteolytically processed into 16 mature nonstructural proteins (Nsps) involved in the assembly of the viral replication-transcription complex (reviewed in (V’Kovski *et al*, 2021)). The Nsps also participate in disrupting host cellular processes and pathways to escape immune recognition and facilitate viral propagation (reviewed in (Minkoff & tenOever, 2023; Suryawanshi *et al*, 2021)).

The maturation step to release the individual Nsp polypeptides is executed by two viral-encoded proteases: Nsp5 (also known as Main Protease, M^Pro^/3C-like protease) and Nsp3 (also known as Papain-Like Protease, PL^Pro^) (Narayanan *et al*, 2022). Nsp3 is contained within pp1a and releases itself from the polyprotein through self-cleavage. Nsp3 also cleaves pp1a at multiple sites to release Nsp1 through Nsp4 (Yan & Wu, 2021). The Nsp5 main protease processes pp1b at eleven sites to release itself and Nsp4 to Nsp16 (Chen *et al*, 2021). Besides cleaving viral substrates, Nsp3 and Nsp5 have also been found to cleave endogenous host proteins linked to the immune response and cell survival (Liu *et al*, 2021; Meyer *et al*, 2021; Meyers *et al*, 2021; Moustaqil *et al*, 2021; Wenzel *et al*, 2021; Zhang *et al*, 2021b). Nsp3 and Nsp5 are essential for viral replication and represent well-characterized drug targets among coronaviruses. Notably, the active component of the antiviral drug Paxlovid is nirmatrelvir, a reversible covalent inhibitor of the Nsp5 main protease (Owen *et al*, 2021). By inhibiting Nsp5 proteolytic activity, Paxlovid reduces viral replication and disease severity in patients with COVID-19.

Purification of individual SARS-CoV-2 proteins from human cells have identified a potential interaction between a catalytic-inactive version of Nsp5 with human tRNA methyltransferase 1 (TRMT1) (Gordon *et al*, 2020b). A cross-coronavirus investigation from the same group has extended this observation by finding that TRMT1 interacts with a catalytic-inactive version of Nsp5 encoded by SARS-CoV-1, which is 98.7% similar to SARS-CoV-2 (Gordon *et al*, 2020a). Interestingly, the same study found no detectable interaction between TRMT1 and Nsp5 encoded by MERS-CoV, which has 66.8% similarity with SARS-CoV-2. TRMT1 was also identified as a potential interactor with catalytic-inactive Nsp5 using proximity-dependent biotinylation (Samavarchi-Tehrani *et al*, 2020).

TRMT1 is a tRNA modification enzyme that catalyzes the formation of dimethylguanosine (m2,2G) at position 26 in more than half of all human tRNAs (Dewe *et al*, 2017; Jonkhout *et al*, 2021). The m2,2G modification is located at a key structural position in tRNAs and is hypothesized to play a role in proper tRNA folding (Pallan *et al*, 2008; Steinberg & Cedergren, 1995). TRMT1-deficient human cells exhibit perturbations in global protein synthesis and decreased proliferation (Dewe *et al*., 2017). Moreover, loss-of-function variants in the *TRMT1* gene are the cause of certain types of intellectual disability disorders in humans (Blaesius *et al*, 2018; Najmabadi *et al*, 2011; Zhang *et al*, 2020). Thus, changes in TRMT1 activity can impact protein synthesis leading to downstream cellular and biological effects.

The life cycle of many viruses has been linked to host tRNA biology (reviewed in (Dremel *et al*, 2023; Nunes *et al*, 2020)). In one well-known example, retroviruses such as HIV use cellular tRNAs as primers for reverse transcription and other viral functions (reviewed in (Jin & Musier-Forsyth, 2019). DNA and RNA viruses can also impact tRNA synthesis, processing, and charging to modulate infection and pathogenesis (Dremel *et al*, 2022; Netzer *et al*, 2009; Tucker *et al*, 2020). In a recent study, Chikungunya RNA virus infection has been shown to increase the expression of a human tRNA modification enzyme and alter tRNA modification patterns to favor viral protein expression (Jungfleisch *et al*, 2022). Interestingly, SARS-CoV-2 viral particles exhibit an enrichment of specific host tRNAs with distinct modification profiles (Pena *et al*, 2022). Moreover, analysis of RNA extracted from human nose swabs have identified tRNA profiles and modification signatures associated with mild versus severe COVID-19 (Katanski *et al*, 2022). The change in tRNA modification profiles could represent another mechanism employed by SARS-CoV-2 to take over translation in addition to previously described strategies (Finkel *et al*, 2021).

The putative interaction between Nsp5 and TRMT1 indicates that SARS-CoV-2 infection could affect the function of TRMT1 with consequent effects on tRNA modification levels. However, the association between Nsp5 and TRMT1 has not been validated nor characterized. Here, we find that TRMT1 is an endogenous cleavage target of Nsp5 resulting in methyltransferase inactive cleavage products. Moreover, we find that SARS-CoV-2 infection correlates with decreased TRMT1 levels and a reduction in m2,2G-modified tRNAs. In addition, we provide evidence that TRMT1 expression is required for efficient SARS-CoV-2 replication in human cells. These studies uncover TRMT1 as a novel proteolytic cleavage target of Nsp5 during SARS-CoV-2 infection and suggest possible pathological mechanisms associated with perturbations in TRMT1-catalyzed tRNA modifications.

## Results

### SARS-CoV-2 infection reduces cellular levels of TRMT1 and TRMT1-catalyzed tRNA modifications

To test the effects of SARS-CoV-2 infection on TRMT1 and tRNA modifications, we used an MRC-5 human lung fibroblast cell line expressing the ACE2 receptor that is permissive for SARS-CoV-2 infection (Raymonda *et al*, 2022; Uemura *et al*, 2021a; Uemura *et al*, 2021b). MRC-5 cells were mock-infected or infected with SARS-CoV-2 followed by harvesting at 24- and 48- hours post-infection for sample preparation. We chose the 24- and 48-hour time points since MRC5-ACE2 cells exhibit elevated accumulation of viral RNA at these time points without extreme cell death that occurs at subsequent time points (Raymonda et al 2022). Immunoblotting for the SARS-CoV-2 nucleocapsid (N) protein confirmed the infection and expression of viral proteins in MRC-5 cells compared to mock infected cells (Figure 1A, N protein, compare lanes 1 and 2 to lanes 3 and 4). To probe for TRMT1, we used an antibody that detects the major TRMT1 isoform of ∼72 kDa as well as a non-specific band at ∼65 kDa in human cell lysates (Dewe *et al*., 2017; Perez *et al*, 2022). Using this antibody, we also detected the 72 kDa TRMT1 isoform as well as the non-specific 65 kDa band in MRC-5 cell lysates (Figure 1A, circle denotes TRMT1, asterisk denotes non-specific band). In multiple independent replicates, MRC-5 cells infected with SARS-CoV-2 exhibited a ∼30% reduction in TRMT1 levels at 24- and 48-hours post-infection compared to mock-infected cells (Figure 1B). These results provide evidence that SARS-CoV-2 infection reduces cellular levels of TRMT1.

**Figure 1.**
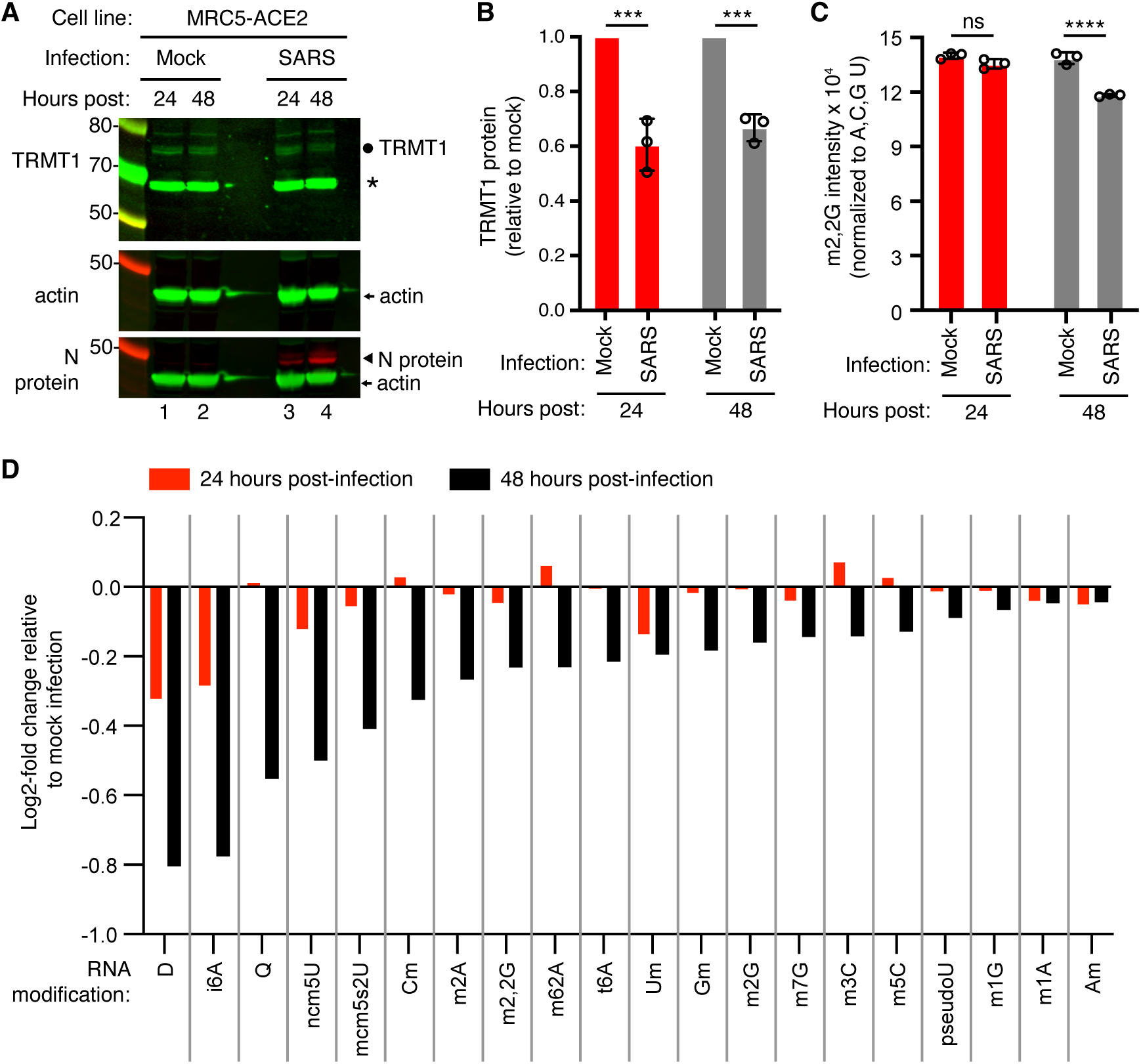
Human cells infected with SARS-CoV-2 exhibit a reduction in TRMT1 levels and perturbations in tRNA modification patterns. (A) Immunoblot analysis of lysates prepared from MRC-5-ACE2 human cells that were mock-infected or infected with SARS-CoV-2 at MOI of 5 for 24 or 48 hours. The immunoblot was probed with anti-TRMT1, actin, or SARS-CoV-2 nucleocapsid (N) antibodies. Circle represents endogenous full-length TRMT1. Asterisk (*) denotes a non-specific band. Size markers are noted in kiloDalton. (B) Quantification of TRMT1 signal intensity normalized to actin in the mock or SARS-CoV-2-infected cell lines. TRMT1 protein levels are expressed relative to mock-infected samples for each time point. (C) m2,2G levels in small RNAs isolated from MRC5 cells that were either mock-infected or infected with SARS-CoV-2 at MOI of 5 for 24 or 48 hours. m2,2G levels were normalized to A, C, G, and U. Samples were measured in biological replicates. Statistical significance for (B) and (C) was determined by two-way ANOVA with multiple comparisons test. ***p<0.001; ****p<0.0001; ns, non-significant. (D) Levels of the indicated RNA modifications in small RNAs isolated from MRC5 cells that were either mock-infected or infected with SARS-CoV-2 for 24 or 48 hours. RNA modification levels were normalized to A, C, G, and U. Y-axis represents the log2 fold change in the levels of the indicated tRNA modification between SARS-CoV-2 infected versus mock-infected MRC5 cells.

We next measured the levels of TRMT1-catalyzed m2,2G modifications in cellular RNA using quantitative mass spectrometry. While m2,2G levels did not exhibit a significant change between mock-infected and SARS-CoV-2-infected cells at 24 hours post-infection, m2,2G levels were decreased by ∼15% at 48 hours post-infection in multiple biological replicates (Figure 1C). The decrease in m2,2G levels after SARS-CoV-2 infection was also reproduced in an independent experiment (Supplemental Figure 1). In addition to m2,2G, more than half of all tested RNA modifications exhibited a decrease in SARS-CoV-2-infected cells compared to mock-infected cells at 24 hours post-infection (Figure 1D). At 48 hours post-infection with SARS-CoV-2, most of the tested RNA modifications exhibited a decrease in levels compared to mock-infected cells. The RNA modifications that decreased upon viral infection include modifications that are known to be present in multiple types of RNA, including tRNA, rRNA, and mRNA. Thus, SARS-CoV-2 infection of human MRC-5 lung fibroblast cells alters the steady-state levels of multiple RNA modifications, including the m2,2G modification catalyzed by TRMT1.

### Nsp5 interacts with TRMT1

Examination of the primary structure of human TRMT1 revealed an eight amino acid residue sequence between the methyltransferase domain and zinc-finger motif matching the consensus sequence of Nsp5 cleavage sites in SARS-CoV-2 polyproteins (Figure 2A, B). The putative cleavage site in TRMT1 includes a glutamine (Q) residue conserved at the fourth position that is found in all Nsp5 cleavage sites of SARS-CoV-2 polyproteins (Figure 2B). Based upon a predicted tertiary structure of TRMT1 using Alpha Fold (Jumper *et al*, 2021; Varadi *et al*, 2022), this putative Nsp5 cleavage site is expected to lie in an unstructured linker region exposed on the surface of TRMT1 (Figure 2C).

**Figure 2.**
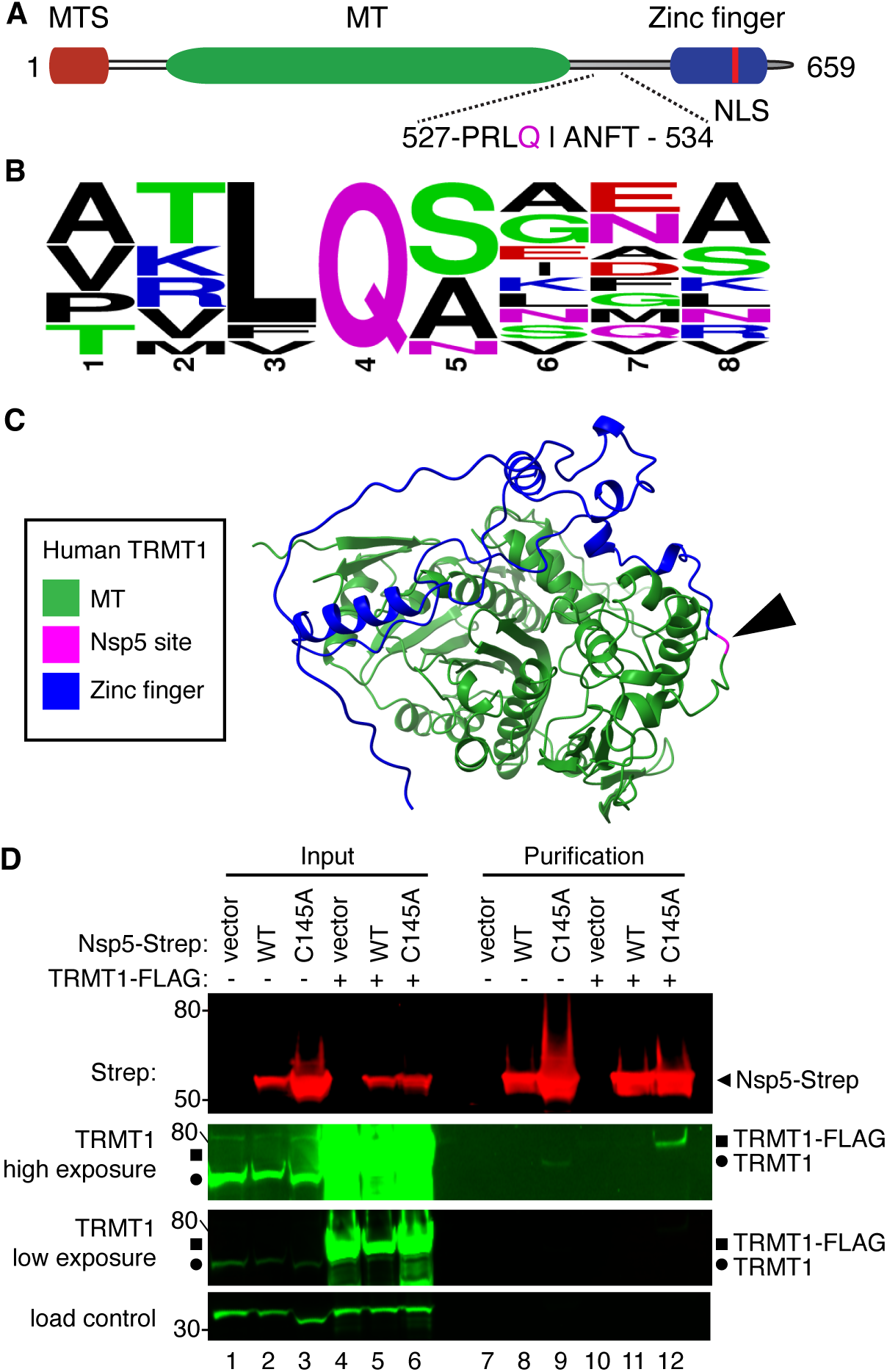
SARS-CoV-2 Nsp5 binds TRMT1 in human cells. (A) Schematic of human TRMT1 primary structure with predicted Nsp5 cleavage site. Mitochondrial targeting signal (MTS), methyltransferase (MT) domain and zinc finger motif are denoted. (B) Consensus sequence logo of cleavage sites in SARS-CoV-2 polyproteins. (C) Alpha-fold predicted structure of human TRMT1 with putative Nsp5 cleavage site denoted in magenta and arrowhead. (D) Immunoblot of input and streptactin purifications from human cells expressing empty vector, wildtype (WT) Nsp5, or Nsp5-C145A fused to the Strep tag without or with co-expression with TRMT1-FLAG. The immunoblot was probed with anti-Strep, FLAG and actin antibodies. Square represents TRMT1-FLAG, circle represents endogenous TRMT1. Size markers are noted to the left in kiloDalton. The experiment in (D) was repeated in Supplemental Figure 2.

To investigate a potential interaction between Nsp5 and TRMT1, we used human 293T cells due to their high transfection efficiency. We transfected human 293T cells with plasmids expressing wildtype (WT) Nsp5 from SARS-CoV-2 or Nsp5-C145A. The Nsp5-C145A mutant contains an alanine substitution of the catalytic cysteine residue in the active site of Nsp5 and is proteolytically inactive (Lee *et al*, 2020). The Nsp5 proteins were expressed as fusion proteins with the Twin-Strep purification tag (Schmidt *et al*, 2013). We also tested the interaction between Strep-tagged Nsp5 or Nsp5-C145A with TRMT1 by co-expression with TRMT1 fused with a carboxyl-terminal FLAG tag. TRMT1 was tagged at the carboxy-terminus to prevent interference with the amino-terminal mitochondrial targeting signal (Dewe *et al*., 2017). The Strep-tagged Nsp5 or Nsp5-C145A mutant were affinity purified from cell lysates on streptactin resin, eluted with biotin and purified proteins detected by immunoblotting. Probing of input lysates and purified samples with anti-Strep antibodies confirmed the expression and recovery of WT-Nsp5 and Nsp5-C145A (Figure 2D, Strep). Nsp5-C145A protein accumulated at higher levels compared to WT-Nsp5 due to the absence of self-cleavage by Nsp5-C145A and the known toxicity of expressing WT-Nsp5 in mammalian cells (Resnick *et al*, 2021; Wenzel *et al*., 2021).

To probe for TRMT1, we used the antibody described above that detects endogenous full-length TRMT1 of 72 kDa (Figure 2D, input lanes 1 to 3, circle denotes endogenous TRMT1). The antibody also detected the overexpressed TRMT1-FLAG protein (Figure 2D, lanes 4 to 6, square denotes TRMT1-FLAG). Only background levels of endogenous TRMT1 or TRMT1-FLAG were detected in the vector or WT-Nsp5 purifications (Figure 2D, TRMT1, low or high exposure, lanes 7, 8, 10 and 11). In contrast, endogenous TRMT1 was enriched in the purification with Nsp5-C145A (Figure 2D, TRMT1 high exposure, lane 9). TRMT1-FLAG was also detected specifically in the Nsp5-C145A purification compared to the control or Nsp5 purifications (Figure 2D, compare lanes 10 and 11 to lane 12). The association between TRMT1 and Nsp5-C145A was reproduced in an independent purification (Supplemental Figure 2). These results provide evidence that Nsp5 can interact with TRMT1 when Nsp5 is in the catalytic-inactive form.

### Nsp5 cleaves TRMT1 in a site-specific manner

The interaction of TRMT1 with Nsp5-C145A along with cleavage site prediction suggests that TRMT1 could be a proteolysis substrate of Nsp5. To test this hypothesis, we monitored for TRMT1 cleavage in 293T human cells expressing Strep-tagged Nsp5 or GFP as a control. Cleavage after amino acid residue 530 of TRMT1 is predicted to result in an N-terminal fragment of 58 kDa and a C-terminal fragment of 14 kDa. To probe for the N-terminal TRMT1 fragment, we used the antibody described above which targets residues 201 to 229 of TRMT1. Using this antibody, we detected full-length TRMT1 and the ∼65 kDa non-specific band in lysates from 293T cells expressing Strep-GFP (Figure 3A, TRMT1, lanes 1 to 3, circle and asterisk, respectively). No detectable change in the levels of full-length TRMT1 was observed in human cells expressing Nsp5 or Nsp5-C145A (Figure 3A, TRMT1, quantified in 3B). However, lysates prepared from human cells expressing Nsp5 exhibited an additional band migrating below the non-specific band that matches the predicted size of the N-terminal TRMT1 fragment (Figure 3B, TRMT1, lanes 4 to 6, arrow). The putative N-terminal TRMT1 fragment was detectable at 24 hours post-transfection with the Nsp5-expression plasmid, increased at 48 hours, and remained detectable at 72 hours post-transfection (Figure 3B). In contrast, the N-terminal TRMT1 fragment was not detected above background signal in human cells expressing the proteolytically inactive Nsp5-C145A mutant (Figure 3B, TRMT1, lanes 7 to 9, quantified in 3C).

**Figure 3.**
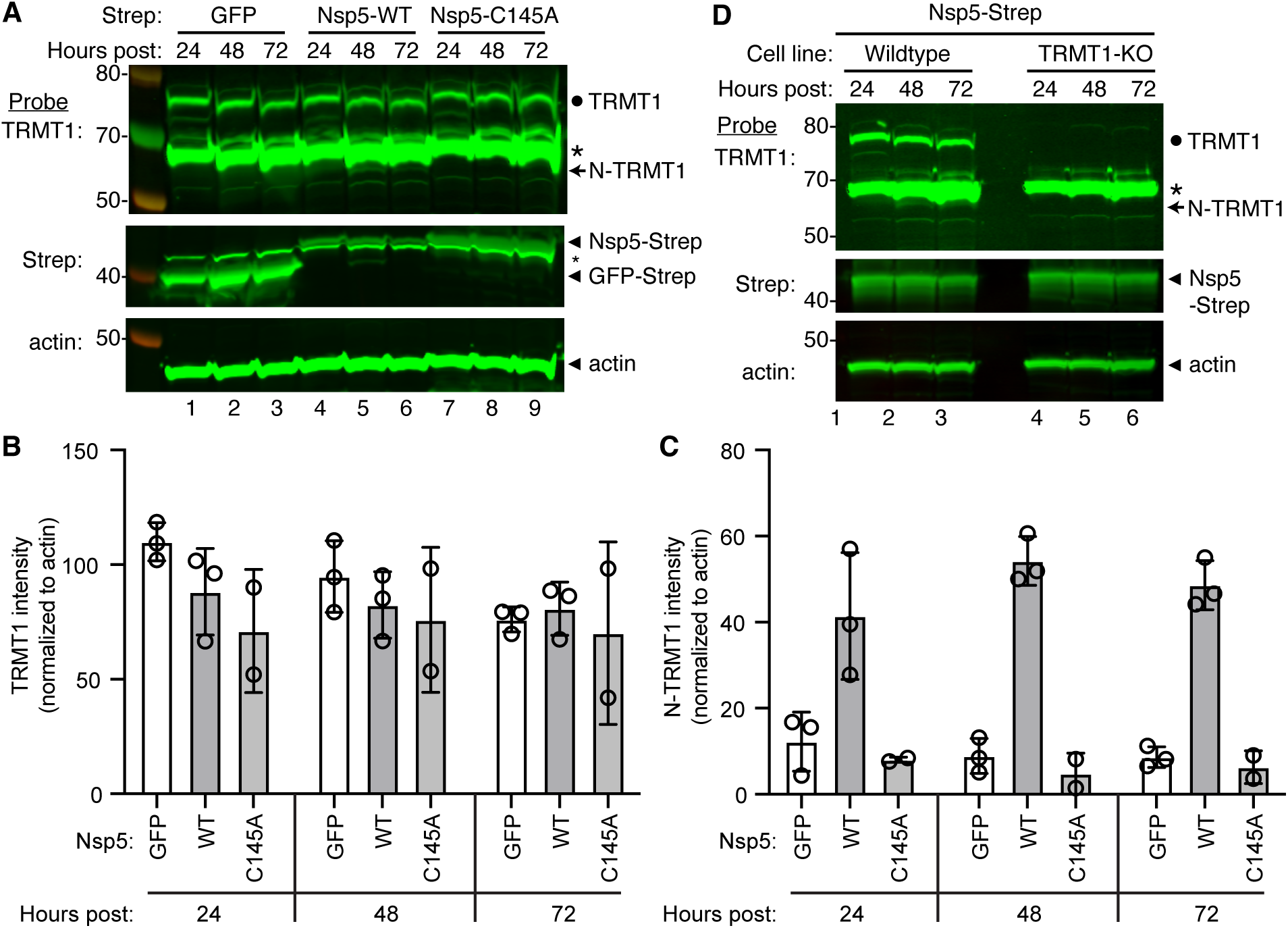
Nsp5 expression induces cleavage of TRMT1 in human cells. (A) Immunoblot of lysates prepared from human 293T cells expressing GFP, Nsp5 or Nsp5-C145A. The immunoblot was probed with anti-TRMT1, Strep, or actin antibodies. Hours post represents the time post-transfection. Circle represents endogenous TRMT1. Arrow represents the N-terminal (N)-TRMT1 cleavage fragment. * denotes a non-specific band. Size markers to the left in kiloDalton. (B, C) Quantification of endogenous TRMT1 or N-terminal (N)-TRMT1 cleavage product in transfected cells. TRMT1 and N-TRMT1 signal was normalized to actin. (D) Immunoblot of lysates prepared from wildtype or TRMT1-knockout (KO) human cell lines expressing Nsp5. Experiments in (A) and (D) were repeated three times (see Source data).

We validated the specificity of our results by using a human 293T TRMT1-knockout (KO) cell line that is deficient in TRMT1 expression (Dewe *et al*., 2017; Zhang *et al*., 2020). In this case, neither full-length TRMT1 nor the TRMT1 cleavage fragment were detected in a TRMT1-deficient human cell line expressing Nsp5 (Figure 3D, compare lanes 1 through 3 to lanes 4 through 6). This result provides additional confirmation that the N-terminal TRMT1 fragment arises from cleavage of endogenous TRMT1 by Nsp5.

We also attempted to detect the C-terminal TRMT1 cleavage fragment in human cells expressing Nsp5 using an antibody targeting residues 609 to 659 of TRMT1 (Zhang *et al*, 2021a). This anti-TRMT1 antibody detects the full-length 72 kDa TRMT1 isoform that is absent in TRMT1-KO cells (Supplemental Figure 3A, lanes 1 and 2). However, no additional band matching the expected size of the C-terminal fragment was detected in human cells expressing Nsp5 compared to cells expressing GFP or Nsp5-C145A (Supplemental Figure 3A, lanes 3 through 6). This could be due to degradation of the C-terminal TRMT1 fragment and/or low sensitivity of the antibody.

To increase the sensitivity for detecting the C-terminal TRMT1 fragment, we co-transfected Nsp5 expression plasmids along with a plasmid encoding TRMT1 fused to a FLAG tag at the C-terminus. Confirming the results above with endogenous TRMT1, the N-terminal TRMT1 cleavage product was detected in the lysate of human cells overexpressing TRMT1-FLAG and Nsp5 but not with vector or Nsp5-C145A (Figure 4A, TRMT1, compare lanes 1 and 3 to lane 2). In addition, using an antibody against the FLAG tag, we could detect a ∼20 kDa product matching the expected molecular weight of a FLAG-tagged C-terminal TRMT1 cleavage product in the lysate of human cells expressing Nsp5 but not vector alone or Nsp5-C145A (Figure 4A, FLAG, arrowhead). To confirm that the 20 kDa band detected with the anti-FLAG antibody was the C-terminal portion of TRMT1, we probed the cell lysates with the anti-TRMT1 antibody targeting residues 609 to 659 noted above. Using this antibody, we detected the same ∼20 kDa band in the lysate of human cells expressing Nsp5 but not vector alone or Nsp5-C145A (Supplemental Figure 3B, lanes 1 to 3). These data provide evidence that SARS-CoV-2 Nsp5 cleaves at the predicted cleavage site in TRMT1 leading to N- and C-terminal fragments in human cells.

**Figure 4.**
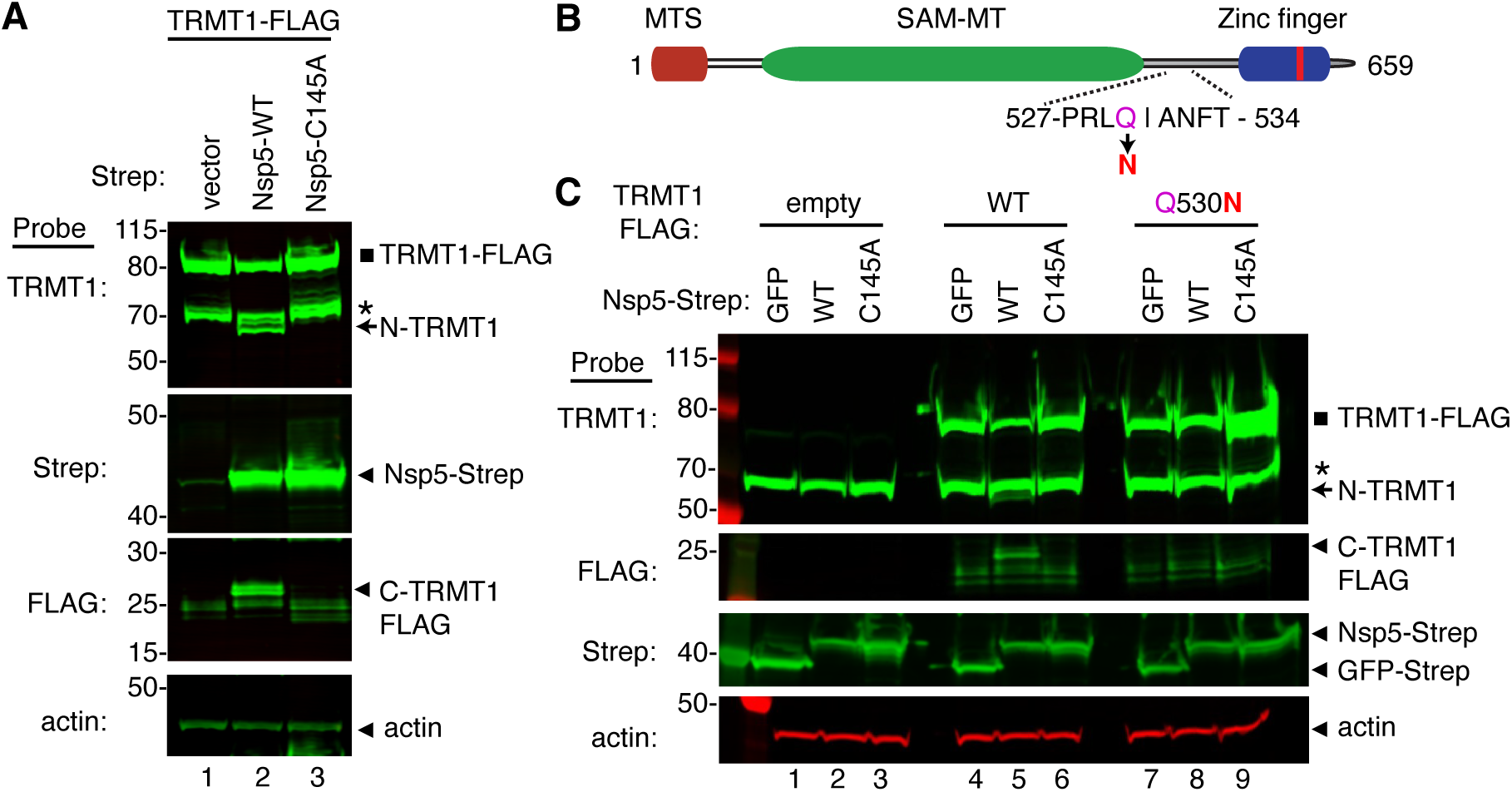
Sequence-dependent cleavage of TRMT1 by SARS-CoV-2 Nsp5. (A) Immunoblot of lysates from human cells expressing empty vector, wildtype (WT) Nsp5-Strep, or Nsp5-C145A-Strep without or with co-expression with TRMT1-FLAG. The immunoblot was probed with anti-Strep, FLAG and actin antibodies. Square represents TRMT1-FLAG, * denotes a non-specific band, arrow represents N-terminal TRMT1 cleavage product and arrowhead indicates the C-terminal TRMT1 cleavage product. (B) Schematic of human TRMT1 with predicted Nsp5 cleavage site and Q530N mutation. (C) Immunoblot of lysates from human cells expressing empty vector, wildtype (WT) Nsp5-Strep, or Nsp5-C145A-Strep without or with co-expression with TRMT1-FLAG or TRMT1-FLAG Q530N. Experiments in (A) and (C) were repeated three times with comparable results (see Source data).

All Nsp5 cleavage sites in SARS-CoV-1 and SARS-CoV-2 polyproteins contain a glutamine residue at position four (Figure 2B)(Grum-Tokars *et al*, 2008; Jin *et al*, 2022; Lee *et al*, 2022). Mutation of the glutamine to asparagine is sufficient to abolish recognition and cleavage by Nsp5 from SARS-CoV-1 or SARS-CoV-2 (Heilmann *et al*, 2022; Muramatsu *et al*, 2013). Thus, we tested if TRMT1 exhibited the same requirements for cleavage by Nsp5 by generating an expression construct for TRMT1-FLAG in which residue Q530 of the predicted Nsp5 cleavage site was mutated to asparagine (Figure 4B, Q530N). Further confirming our results above, expression of TRMT1-FLAG with WT-Nsp5 but not GFP or Nsp5-C145A led to the accumulation of N- and C-terminal TRMT1 cleavage fragments (Figure 4C, TRMT1 and FLAG, lanes 4 to 6, arrow and arrowhead, Supplemental Figure 3B). In contrast to wildtype TRMT1, the appearance of the N- or C-terminal TRMT1 cleavage fragments was barely detectable when the TRMT1-Q530N mutant was co-expressed with Nsp5 (Figure 4C, lane 8, Supplemental Figure 3B). Altogether, these results demonstrate that TRMT1 can be recognized and cleaved by Nsp5 in human cells with cleavage requiring a sequence that matches Nsp5 cleavage sites in SARS-CoV-2 polyproteins.

### Functional properties of TRMT1 fragments resulting from Nsp5 cleavage

Cleavage of TRMT1 after residue Q530 will result in an N-terminal fragment encompassing the methyltransferase domain and a C-terminal TRMT1 fragment containing a zinc finger motif that mediates tRNA interaction (Dewe *et al*., 2017; Zhang *et al*., 2020). Thus, we tested the functional properties of the predicted TRMT1 cleavage fragments compared to full-length TRMT1. First, we tested the interaction between the TRMT1 cleavage fragments and RNA. We have previously shown that TRMT1 exhibits a stable interaction with rRNAs and substrate tRNAs that are targets for m2,2G modification (Dewe *et al*., 2017; Zhang *et al*., 2020). Based upon this interaction, we expressed full-length TRMT1, the TRMT1-Q530N mutant, or the TRMT1 fragments as FLAG-tagged fusion proteins in 293T human cells followed by affinity purification and analysis of copurifying RNAs (Figure 5A). Immunoblotting confirmed the expression and purification of each TRMT1 variant on anti-FLAG resin (Figure 5B). As expected, the purification of full-length TRMT1 resulted in the enrichment of rRNA and tRNAs compared to the control purification from vector-transfected cells (Figure 5C, compare lane 6 to lane 7). The Q530N variant of TRMT1 also exhibited comparable binding of tRNAs as full-length TRMT1 (Figure 5C, lane 10). In contrast, no detectable enrichment of rRNA or tRNA was detected for the N-terminal TRMT1 fragment (Figure 5C, lane 8). Interestingly, the TRMT1 C-terminal fragment exhibited similar levels of tRNA binding as full-length TRMT1 (Figure 5C, compare lane 7 to 9). Thus, the N-terminal TRMT1 cleavage fragment appears to be insufficient for binding tRNA while the C-terminal fragment containing the Zn-finger motif is sufficient for binding to tRNAs.

**Figure 5.**
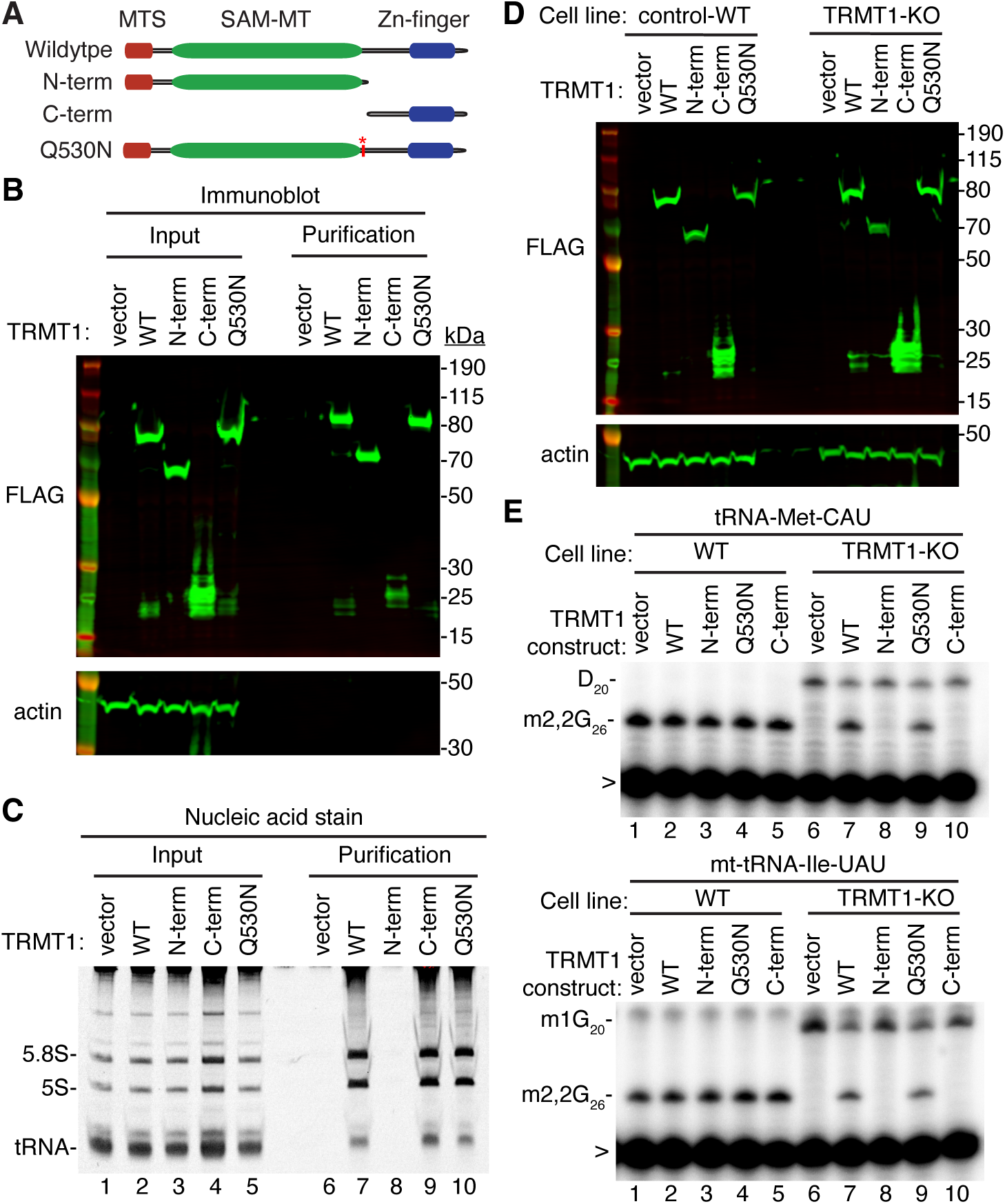
N- and C-terminal TRMT1 cleavage fragments exhibit alterations in RNA binding and tRNA modification activity. (A) Schematic of wildtype TRMT1 and predicted TRMT1 fragments resulting from Nsp5 cleavage at Q530N. (B) Immunoblot analysis of anti-FLAG purifications from human cells expressing vector control, full-length TRMT1, or TRMT1 cleavage fragments fused to the FLAG tag. The immunoblot was probed with anti-FLAG and anti-actin antibodies. (C) Nucleic acid stain of RNAs extracted from the indicated input or purified samples after denaturing PAGE. The migration pattern of 5.8S rRNA (∼150 nt), 5S rRNA (∼120 nt) and tRNAs (∼70–80 nt) are denoted. (D) Immunoblot of TRMT1 expression in either control-WT or TRMT1-KO human 293T cell lines. (E, F) Representative gel of primer extension assays to monitor the presence of m2,2G in tRNA-Met-CAU or mt-tRNA-Ile-UAU from the cell lines transfected with the indicated TRMT1 constructs. D, dihydrouridine; m1G, 1-methylguanosine; >, labeled oligonucleotide used for primer extension. Protein-RNA purification was repeated with comparable results (see Source data for repeat).

To further dissect the functionality of the TRMT1 cleavage fragments, we next used the TRMT1-KO human cell line described above. This TRMT1-KO cell line is deficient in TRMT1 and lacks m2,2G modifications in all tested tRNAs containing G at position 26 (Dewe *et al*., 2017; Zhang *et al*., 2020). Using transient transfection, we expressed full-length TRMT1 or the TRMT1 variants in either the control wildtype (WT) or TRMT1-KO cell lines (Figure 5D). We then assessed for rescue of m2,2G formation in nuclear-encoded tRNA-Met-CAU or mitochondrial-encoded (mt)-tRNA-Ile-UAU using a reverse transcriptase (RT)-based primer extension assay. Based upon this assay, vector-transfected WT human cells exhibited an RT block at position 26 of tRNA-Met-CAU and mt-tRNA-Ile-UAU indicative of the m2,2G modification (Figure 5E, Lane 1). No read-through product was detected for either tRNA in control human cells indicating that nearly all endogenous tRNA-Met-CAU and mt-tRNA-Ile-UAU is modified with m2,2G. Consistent with this observation, increased expression of full-length TRMT1 or variants in control 239T cells had no detectable effect on m2,2G modification in tRNA-Met-CAU or mt-tRNA-Ile-UAU (Figure 5E, lanes 2 through 5).

As expected, the m2,2G modification was absent in tRNA-Met-CAU and mt-tRNA-Ile-UAU isolated from the vector-transfected TRMT1-KO cell line leading to read-through to the next RT block (Figure 5E, lane 6). Re-expression of full-length TRMT1 or TRMT1-Q530N in the TRMT1-KO cell line was able to restore m2,2G formation in both tRNA-Met-CAU and mt-tRNA-Ile-UAU (Figure 5E, lanes 7 and 9). However, neither the N- nor C-terminal TRMT1 fragment was able to restore m2,2G formation in the tRNAs of the TRMT1-KO cell line (Figure 5E, lanes 8 and 10). These results indicate that cleavage of TRMT1 by Nsp5 leads to protein fragments that are inactive for tRNA modification activity.

In addition to TRMT1 activity, we compared the subcellular localization pattern of full-length TRMT1 to the TRMT1 cleavage fragments. To visualize TRMT1, we transiently transfected 293T human cells with constructs expressing human TRMT1 fused at its carboxy terminus to GFP. As a mitochondria marker, we co-expressed a red fluorescent protein fused to the mitochondria targeting signal of pyruvate dehydrogenase (Supplemental Figure 4, Mito-RFP). Full-length TRMT1-GFP exhibited cytoplasmic localization along with a punctate signal in the nucleus (Supplemental Figure 4, TRMT1-WT, GFP). The fluorescence signal for TRMT1 in the cytoplasm overlapped partially with the RFP-tagged mitochondrial marker (Supplemental Figure 4, Merge, yellow signal). The mitochondrial and nuclear localization pattern of full-length TRMT1 is consistent with prior studies demonstrating the presence of a mitochondrial targeting signal and nuclear localization signal in TRMT1 (Dewe *et al*., 2017). The N-terminal TRMT1 fragment exhibited primarily cytoplasmic localization with greatly reduced signal in the nucleus, consistent with lack of the nuclear localization signal at the C-terminus (Supplemental Figure 4, TRMT1 N-term, GFP, merge). In contrast to the N-terminal TRMT1 fragment, the C-terminal TRMT1 fragment displayed primarily nuclear localization with very little signal in the cytoplasm (Supplemental Figure 4, TRMT1 C-term, GFP, merge). These results suggest that TRMT1 in the cytoplasm is the likely target of Nsp5 cleavage with the cleavage fragments displaying altered localization compared to full-length TRMT1.

### TRMT1-deficient human cells exhibit reduced levels of SARS-CoV-2 RNA replication

We next investigated whether TRMT1 expression impacts SARS-CoV-2 replication by infecting the 293T control-wildtype (WT) or TRMT1-KO human cell lines described above with SARS-CoV-2. To render the 293T cell lines permissive for SARS-CoV-2 infection, the cell lines were engineered to stably express the ACE2 receptor from an integrated lentiviral vector (Supplemental Figure 5A). Immunoblotting for the SARS-CoV-2 nucleocapsid protein confirmed that SARS-CoV-2 could infect both the control-WT and TRMT1-KO human cell lines expressing ACE2 (Supplemental Figure 5B). Using these cell lines, we first monitored the effect of SARS-CoV-2 infection on TRMT1 levels. As expected, there was no detectable TRMT1 in any of the lanes containing lysate from the TRMT1-KO cell line (Figure 6A, compare control-WT, lanes 1 to 3, to TRMT1-KO, lanes 4 to 6). In the control-WT cells, we detected a decrease in TRMT1 levels at both multiplicity of infection (MOI) of 0.2 and 0.4 (Figure 6A, TRMT1, quantified in Figure 6B). The decrease in TRMT1 in SARS-CoV-2-infected 293T cells is comparable to the reduction in TRMT1 levels in MRC5-ACE2 human cell lines infected with SARS-CoV-2 (Figure 1). We also compared the levels of endogenous TRMT1 after infection with SARS-CoV-2 at a higher MOI of 5.0. Human 293T cells infected at the higher MOI exhibited a reduction in TRMT1 levels to nearly the background of the TRMT1-KO cell lines (Figure 6C and D). These results indicate that SARS-CoV-2 infection can induce a drastic reduction in cellular TRMT1 levels.

**Figure 6.**
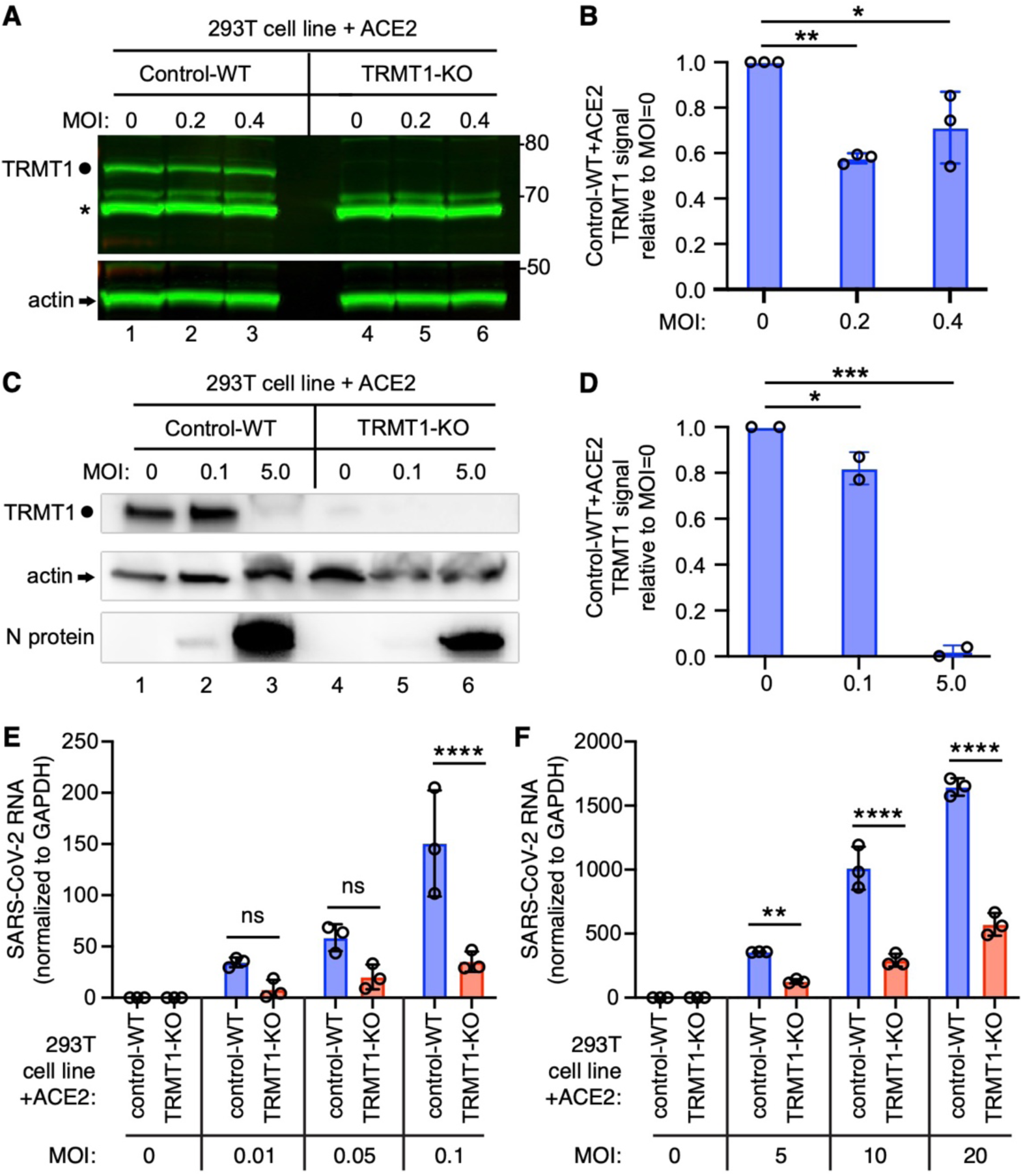
The expression of TRMT1 affects the levels of SARS-CoV-2 RNA replication in human cells. (A) Immunoblot of lysates prepared from 293T control-wildtype (WT) or TRMT1-KO cell lines that were mock-infected (MOI of 0) or infected with SARS-CoV-2 at MOI of 0.2 or 0.4 for 24 hours. The immunoblot was probed with antibodies against TRMT1 or actin. Circle represents endogenous full-length TRMT1. Asterisk denotes a non-specific band. Size markers to the right in kiloDalton. (B) Normalized TRMT1 signal intensity relative to mock-infected cells (MOI of 0). (C) Immunoblot of lysates prepared from 293T control-wildtype (WT) or TRMT1-KO cell lines that were mock-infected (MOI of 0) or infected with SARS-CoV-2 at MOI of 0.1 or 5.0 for 24 hours. The immunoblot was probed with antibodies as in (A). (D) Normalized TRMT1 signal intensity relative to mock-infected cells (MOI of 0). (E, F) SARS-CoV-2 RNA copy number in control-WT or TRMT1-KO human 293T cell lines after infection at the indicated MOI for 24 hours. Viral copy number was measured by QRT-PCR and normalized to GAPDH. Samples were measured in triplicate. *p < 0.05; **p<0.01; ***p<0.001; ****p < 0.0001; ns, non-significant.

The ACE2-expressing cell lines were then infected at low or high multiplicity of infection (MOI) followed by harvesting of cells at 24 hours post-infection. Intracellular viral RNA levels were monitored by quantitative RT-PCR of the SARS-CoV-2 envelope protein gene. As expected, titration of SARS-CoV-2 viral particles at either low or high MOI led to a concomitant increase in viral RNA in both control and TRMT1-KO cell lines (Figure 6E and F). Notably, TRMT1-KO cells exhibited a ∼3 to 4-fold reduction in viral RNA compared to control cells infected at the same MOI (Figure 6E and F). These results suggest that expression of TRMT1 is necessary for efficient SARS-CoV-2 replication in human cells.

To confirm that TRMT1-deficiency is the cause of the reduced SARS-CoV-2 replication, we generated TRMT1-KO cell lines re-expressing TRMT1-WT or TRMT1-Q530N with a C-terminal FLAG tag from an integrated lentiviral construct. As expected, wildtype control 293T cells expressed endogenous TRMT1 that was absent in the TRMT1-KO cell lines with empty vector (Supplemental Figure 6, TRMT1, lanes 1 and 2). Furthermore, we could detect re-expression of TRMT1-WT or TRMT1-Q530N in the TRMT1-KO cell lines containing the integrated lentiviral TRMT1 expression vectors (Supplemental Figure 6, TRMT1, lanes 3 to 6). We also validated the cell lines by checking for cleavage of the re-expressed TRMT1 by Nsp5. Indeed, the N-terminal TRMT1 cleavage product was detected in the lysate of TRMT1-KO cells expressing TRMT1-WT and Nsp5, but not TRMT1-Q530N and Nsp5 (Supplemental Figure 6, TRMT1, compare lanes 3 and 4). These results further support our findings described above that Nsp5 expression leads to site-specific cleavage of TRMT1 in human cells.

We then expressed the ACE2 receptor in the complemented TRMT1-KO cell lines to render them permissive for SARS-CoV-2 infection (Supplemental Figure 7). Using these cell lines, we first tested the effect of SARS-CoV-2 infection on TRMT1 levels. Compared to mock-infection, TRMT1 protein levels decreased in the TRMT1-KO cell lines expressing TRMT1-WT after infection with SARS-CoV-2 (Figure 7A, lanes 1 to 3; quantified in 7B). Notably, the level of TRMT1 was not appreciably changed in the TRMT1-KO cell line expressing TRMT1-Q530N after infection with SARS-CoV-2 (Figure 7A, lanes 4 to 6; quantified in 7C). These results provide evidence that the decrease in endogenous TRMT1 levels after SARS-CoV-2 infection is due to cleavage of TRMT1 by Nsp5.

**Figure 7.**
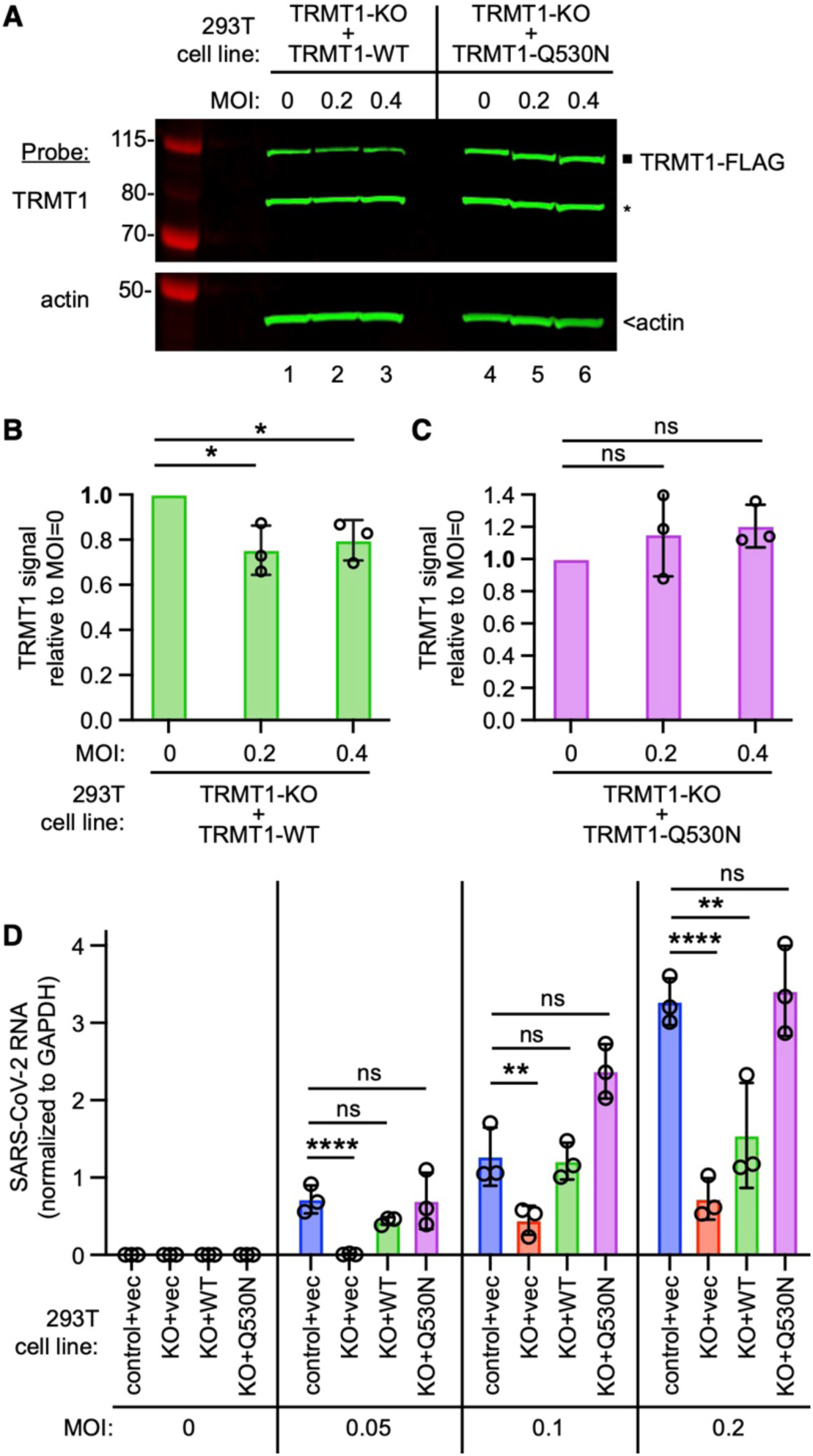
TRMT1 is required for efficient SARS-CoV-2 replication in human cells. (A) Immunoblot of lysates prepared from the indicated 293T TRMT1-KO cell lines that were mock-infected (MOI of 0) or infected with SARS-CoV-2 for 24 hours. The immunoblot was probed with antibodies against TRMT1 or actin. Square represents full-length TRMT1-FLAG. Asterisk denotes a non-specific band. Size markers to the left in kiloDalton. (B) Normalized TRMT1-WT signal intensity relative to mock-infected cells (MOI of 0). (C) Normalized TRMT1-Q530N signal intensity relative to mock-infected cells (MOI of 0). (D) SARS-CoV-2 RNA copy number in control-WT or TRMT1-KO human 293T cell lines after infection at the indicated MOI. Viral copy number was measured by QRT-PCR and normalized to GAPDH. *p < 0.05; **p<0.01; ****p < 0.0001; ns, non-significant.

We next measured SARS-CoV-2 RNA levels after infecting the TRMT1-KO cell lines with SARS-CoV-2. Reproducing our results above, we found that the TRMT1-KO+vector cell line exhibited a decrease in viral RNA levels compared to control cells infected at the same MOI (Figure 7D, compare control+vec to KO+vec). TRMT1-KO cell lines re-expressing TRMT1 exhibited higher viral RNA production relative to the TRMT1-KO+vector cell line that was comparable to the control-WT cells (Figure 7D, compare KO+vec to KO+WT). Notably, the TRMT1-KO cell line expressing TRMT1-Q530N exhibited a higher level of viral RNA production compared to the TRMT1-KO cell line expressing TRMT1-WT at higher MOI (Figure 7, KO+Q530N). Altogether, these results uncover a requirement for TRMT1 expression for efficient SARS-CoV-2 replication in human cells.

### Impact of TRMT1 on SARS-CoV-2 particle infectivity

Since the presence or absence of TRMT1 appears to impact SARS-CoV-2 RNA production in human 293T cells, we tested if this phenomenon could alter viral particle production. To this end, we collected tissue culture media supernatant containing viral particles generated from control-WT or TRMT1-KO cell lines that were infected by SARS-CoV-2 for 24 hours. The viral titer of the supernatant was determined by focus forming unit assay and particle infectivity expressed relative to viral genomic RNA copy number within the same sample. Consistent with the reduced levels of viral RNA in TRMT1-KO cells infected with SARS-CoV-2, the supernatant from infected TRMT1-KO cells exhibited reduced titer and slightly lower infectivity when compared to viral particles produced from control-WT cells (Figure 8, control-WT versus TRMT1-KO). Re-expression of TRMT1 in the TRMT1-KO cells partially restored viral titer at the lower MOI and increased infectivity compared to supernatants from TRMT1-KO cell lines (Figure 8, TRMT1-KO+vec versus TRMT1-KO+WT). Interestingly, the expression of the non-cleavable TRMT1-Q530N variant in TRMT1-KO cells promoted an increase of viral titer as well as infectivity compared to expression of wildtype TRMT1 (Figure 8, TRMT1-KO+WT versus TRMT1-KO+Q530N). Altogether, these observations suggest an unexpected role for TRMT1 expression in virus production and the generation of optimally infectious SARS-CoV-2 particles.

**Figure 8.**
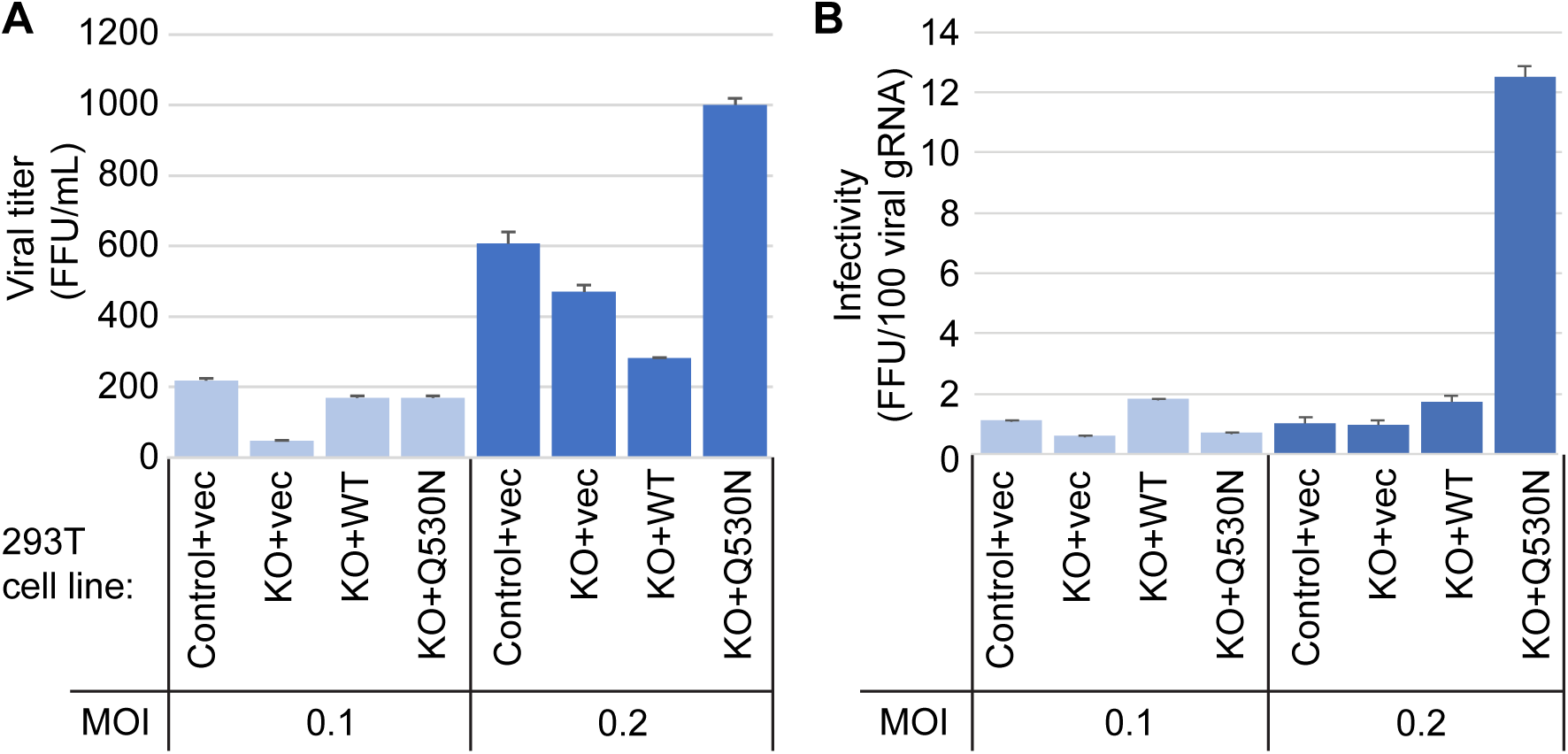
Viral infectivity measurement of supernatants collected from the indicated cell lines infected with SARS-CoV-2 for 24 hours. (A) Viral titer of supernatants collected from the indicated cell lines infected with SARS-CoV-2. Infectious titer was determined by TCID50 endpoint dilution assay in VeroE6 cells and expressed in focus forming units per mL of supernatant (FFU/mL). (B) Infectivity of SARS-CoV-2 particles generated from cell lines in (A). Infectivity of viral particles was calculated with the formula [(FFU/mL)/(viral genomic RNA copies/mL)], and expressed in FFU per 100 genomic copies.

## Discussion

Here, we demonstrate that TRMT1 tRNA modification enzyme is an endogenous cleavage target of the SARS-CoV-2 main protease. Our studies parallel the simultaneous work by D’Oliviera *et al*. that has elucidated how the SARS-CoV-2 main protease recognizes the TRMT1 cleavage sequence (D’Oliviera *et al*, 2023; D′Oliviera *et al*, 2023). D’Oliviera *et al*. have determined the structural basis for recognition of TRMT1 by the SARS-CoV-2 main protease, which has revealed a distinct binding mode for certain substrates of the main protease, including TRMT1. Together, our study and the investigation by D’Oliviera *et al*. provide independent corroboration of each other’s main conclusion that TRMT1 is recognized and cleaved by the SARS-CoV-2 main protease in a sequence-specific manner. Moreover, after our manuscript and the manuscript by D’Oliviera *et al*. were deposited on *BioRxiv*, an independent study by Lu and Zhou has further confirmed our findings that the SARS-CoV-2 main protease can cleave TRMT1 (Lu & Zhou, 2023).

TRMT1 could represent a coincidental substrate of Nsp5 during SARS-CoV-2 infection due to the presence of a sequence matching the cleavage site of SARS-CoV-2 polyproteins. Since the majority of TRMT1 exhibits steady-state localization to the nucleus and mitochondria of multiple human cell types (Dewe *et al*., 2017), Nsp5 could have access mainly to newly synthesized TRMT1 in the cytoplasm. Moreover, the effect of SARS-CoV-2 infection on TRMT1 levels could depend on the concentration of TRMT1, half-life of TRMT1, and/or amount of Nsp5 expression in the infected cell. This could account for the observation that TRMT1 levels are reduced but not abolished in human cells infected with SARS-CoV-2. Consistent with our results, a subset of proteomic studies has found decreased TRMT1 protein levels in SARS-CoV-2-infected human cells as well as in post-mortem tissue samples from deceased COVID-19 patients (Bojkova *et al*, 2020; Nie *et al*, 2021). These findings suggest that SARS-CoV-2 infection could impact TRMT1 protein levels and tRNA modification patterns within the cells of an infected individual.

While TRMT1 cleavage could be a collateral effect of SARS-CoV-2 infection, the subsequent impact on tRNA modification levels could have physiological consequences on downstream molecular processes that ultimately affect cellular health and/or viral replication. We have previously shown that TRMT1-deficient human cells exhibit decreased levels of global protein synthesis, perturbations in redox metabolism, and reduced proliferation (Dewe *et al*., 2017). Moreover, we have found that partial depletion of TRMT1 is sufficient to increase the oxidative stress sensitivity of human neural stem cells. Thus, the reduction in TRMT1 and TRMT1-catalyzed tRNA modifications observed in human lung cells upon SARS-CoV-2 infection could lead to changes in protein synthesis that affects cellular proliferation and metabolism. In lung tissues, the disruption of TRMT1-dependent processes caused by changes in TRMT1-catalyzed tRNA modification levels could contribute to the pathophysiological outcomes associated with COVID-19 disease. Consistent with this possibility, TRMT1 has been identified as a prognosis factor for SARS-CoV-2 disease severity (Li *et al*, 2021a).

We have found that human cells deficient in TRMT1 display reduced viral RNA levels compared to wildtype cells after SARS-CoV-2 infection. This finding suggests that TRMT1 expression is required for efficient SARS-CoV-2 replication. As mentioned above, TRMT1-deficient human cells exhibit an overall reduction in global protein synthesis due to the loss of m2,2G modifications in tRNAs (Dewe *et al*., 2017). Thus, the TRMT1-deficient human cells could present a cellular environment that is unable to support the levels of translation necessary for efficient virus production. In addition, TRMT1-deficient human cells could exhibit changes in gene expression and cellular metabolism that are suboptimal for SARS-CoV-2 replication.

While TRMT1 could be an unintentional target of Nsp5, it remains conceivable that TRMT1 cleavage modulates the SARS-CoV-2 life cycle through an uncharacterized process. One possibility is that viral RNAs could be substrates of TRMT1-catalyzed RNA methylation. Previous studies have found that endogenous host RNA modification enzymes can modify the genomic and sub-genomic RNAs of SARS-CoV-2 (Burgess *et al*, 2021; Di Giorgio *et al*, 2020; Li *et al*, 2021b; Peng *et al*, 2022; Zhang *et al*, 2021c). Moreover, uncharacterized modifications have been identified through Nanopore sequencing in the genomic and subgenomic RNAs of SARS-CoV-2 (Kim *et al*, 2020). There could be portions of the SARS-CoV-2 genome or subgenomic RNAs that fold into substrates that resemble tRNA targets of TRMT1. There has been precedence for the folding of certain segments of plant viral RNAs into tRNA-like structures that can be modified by cellular tRNA modification enzymes (Baumstark & Ahlquist, 2001; Becker *et al*, 1998; Lesiewicz & Dudock, 1978). Moreover, the N-terminal TRMT1 fragment could gain the ability to modify viral RNAs due to altered binding specificity since the N-terminal TRMT1 fragment retains the entire methyltransferase domain but not the Zn-finger motif involved in tRNA interaction. Future studies will examine for possible interactions between TRMT1 and the SARS-CoV-2 transcriptome as well as the presence of TRMT1-catalyzed modifications in SARS-CoV-2 RNAs.

We have previously found that TRMT1-deficient human cells exhibit nearly undetectable m2,2G without major impact on other RNA modifications (Dewe *et al*., 2017). In contrast, human cells infected with SARS-CoV-2 exhibited a decrease in multiple RNA modifications present in a variety of RNAs, including tRNA, rRNA, snRNA, and mRNA. Thus, the widespread changes in RNA modifications after infection with SARS-CoV-2 suggest that coronavirus infection could alter the levels or activity of multiple RNA modification enzymes in addition to TRMT1. Consistent with this hypothesis, RNA modification profiles are changed in response to various forms of cellular stress, including viral infection (Chan *et al*, 2018; Jungfleisch *et al*., 2022). Overall, the study presented here highlights the expanding role of RNA modifications in modulating cellular responses to pathogens that will be the important subject of further investigation.

### Ideas and Speculation

Our findings suggest that the SARS-CoV-2 might self-limit its replication by altering the host translation machinery through TRMT1 degradation and reduced levels of m2,2G-modified tRNAs. This perturbation of the tRNA pool may further inhibit host translation that is already targeted by Nsp1 blockade of mRNA entry on 40S ribosomes (Kamitani *et al*, 2009; Lapointe *et al*, 2021; Schubert *et al*, 2020; Thoms *et al*, 2020; Tidu *et al*, 2020). The inhibition of host translation may be beneficial to certain viruses that can locally maintain a tRNA pool optimized for viral translation (Hernandez-Alias *et al*, 2021; Pavon-Eternod *et al*, 2013; van Weringh *et al*, 2011; Yang *et al*, 2021). Another possibility is that Nsp5-TRMT1 interaction facilitates the packaging of specific tRNAs into viral particles as suggested previously (Pena *et al*., 2022). The observation that expression of the non-cleavable TRMT1-Q530N variant enhances viral replication and infectivity supports the hypothesis that TRMT1 could facilitate tRNA uptake into viral particles. The packaging of specific tRNAs in viral particles could augment viral translation in the subsequent round of infection, thereby enhancing infectivity and perhaps facilitating the species jump of SARS-CoV-2 towards hosts with incompatible codon bias.

## Materials and Methods

### Cell culture

293T human embryonic cell lines were cultured in Dulbecco’s Minimal Essential Medium (DMEM) containing 10% fetal bovine serum, 2 mM L-alanyl-L-glutamine (GlutaMax, Gibco) and 1% Penicillin/Streptomycin. Cells were grown at 37 °C, 20% Oxygen, and 5% CO2. Telomerase-immortalized MRC5 fibroblasts expressing ACE2 (MRC5-ACE2 cells) were cultured in Dulbecco’s modified Eagle serum (DMEM; Invitrogen) supplemented with 10% (vol/vol) fetal bovine serum (FBS) (Atlanta Biologicals), 4.5 g/L glucose, and 1% penicillin-streptomycin (Pen-Strep; Life Technologies) at 37 °C in a 5% (vol/vol) CO2 atmosphere. Vero-E6 cells were cultured in Eagle’s Minimum Essential Medium (ATCC, #30–2003) supplemented with 10% (vol/vol) fetal bovine serum (FBS) (Atlanta Biologicals), 4.5 g/L glucose, 1X Glutamax (Life Technologies, #35050061), and 1% penicillin-streptomycin (Pen-Strep; Life Technologies, #15140122) at 37 °C in a 5% (vol/vol) CO2 atmosphere.

### SARS-CoV-2 infection of human cells for protein and RNA analysis

The SARS-CoV-2 isolate, Hong Kong/VM20001061/2020, was previously isolated from a nasopharyngeal aspirate and throat swab from an adult male patient in Hong Kong and was obtained through BEI resources (NR-52282). Viral stocks of SARS-CoV-2 were propagated in Vero-E6 cells in MEM supplemented with 2% (vol/vol) FBS, 4.5 g/L glucose, 1X Glutamax and 1% penicillin-streptomycin at 37 °C. Viral stock titers were determined by TCID50 analysis in Vero-E6 cells. Experiments involving live SARS-CoV-2 were conducted in a biosafety level 3 facility at the University of Rochester.

For infection, MRC5-ACE2 cells were grown in 6-well plates to 90% confluence in growth medium (MRC5-ACE2: DMEM supplemented with 10% FBS and 1% Pen-Strep). Prior to infection, cells were washed with 1 mL of DMEM supplemented with 2% FBS and 1 % Pen-Strep. For infection, 750 μL of viral master mix (MOI = 5) was added to each well for 1.5 hr. After the adsorption period, medium was removed and replaced with fresh DMEM supplemented with 2% FBS and 1% Pen-Strep. To harvest protein, the cell monolayer was washed with 1 mL cold PBS and cells were scraped into 250 μL disruption buffer (250 mM Tris-HCl Ph 7.4, 10% glycerol, 2% β-mercaptoethanol, 5% SDS). DNA was sheared using a sonication probe and samples were stored at −20 °C. To harvest RNA, medium was removed, and the monolayer was washed with PBS. Cells were collected in 400 μL Trizol and stored at −80 °C.

### Liquid chromatography-mass spectrometry of nucleosides

Total RNA was isolated using Trizol RNA extraction. Small RNAs were subsequently purified from total RNA using the Zymo RNA Clean & Concentrator-5 kit. Small RNAs (1 μg) was digested and analyzed by LC-MS as previously described (Dewe *et al*., 2017; Zhang *et al*., 2020) (Cai *et al*, 2015). Briefly, ribonucleosides were separated using a Hypersil GOLD C18 Selectivity Column (Thermo Scientific) followed by nucleoside analysis using a Q Exactive Plus Hybrid Quadrupole-Orbitrap. The modification difference ratio was calculated using the m/z intensity values of each modified nucleoside following normalization to the sum of intensity values for the canonical ribonucleosides; A, U, G and C. Statistical analysis of the mass spectrometry results was performed using GraphPad Prism software with error bars representing the standard deviation. Statistical tests and the number of times each experiment was repeated are noted in the figure legend. Raw intensity values for each measured nucleoside are provided in the source data file.

### In silico analysis of TRMT1 structure

The Nsp5 cleavage site sequence logo was generated using: https://weblogo.berkeley.edu/logo.cgi

The predicted tertiary structure of human TRMT1 (Uniprot Q9NXH9) was determined using AlphaFold. Structural visualization was performed using UCSF Chimera software developed by the Resource for Biocomputing, Visualization, and Informatics at the University of California, San Francisco (Pettersen *et al*, 2004). The PDB file for the predicted TRMT1 structures has been provided in the source file.

### Plasmid constructs

The pcDNA3.1-TRMT1-FLAG and pcDNA3.1-TRMT1-GFP expression plasmids have been described previously (Dewe *et al*., 2017). The pcDNA3.1-TRMT1-FLAG-Q530N expression construct was generated by DpnI site-directed mutagenesis. The pcDNA3.1 expression plasmids encoding the N- and C-terminal TRMT1 fragments were generated by PCR cloning. Lentiviral constructs expressing TRMT1-WT or TRMT1-Q530N were generated by T5 Exonuclease DNA Assembly (TEDA) of PCR fragments into pLenti-CMV-GFP-Blast (Addgene 17445) (Xia *et al*, 2019). All plasmid constructs were verified by Sanger sequencing and whole plasmid sequencing (Plasmidsaurus).

### Generation of human cell lines by lentiviral integration

For lentivirus production, 1 x 10^6^ 293T cells were seeded onto 60 mm tissue culture dishes. After 24 hours, 1.5 μg of pLenti CMV Blast plasmids containing empty vector or TRMT1 along with 1.5 μg of psPAX2 packaging plasmid and 0.7 μg of pMD2.G envelope plasmid was transfected into the 293T cells using calcium phosphate transfection. Media was changed 16 hours post-transfection. At 48 hours post-transfection, the media containing virus was collected, filter sterilized through a 0.45 μm filter, flash frozen, and stored at −80 °C.

For lentiviral infection of 293T cell lines, 1 x 10^6^ cells were seeded onto 60 mm tissue culture dishes. After 24 hours, 1 mL of either virus or media for mock infection along with 2 mL of media supplemented with 10 μg/mL of polybrene was added to each well. The cells were washed with PBS and fed fresh media 24 hours post infection. Polyclonal cell lines with stable integration of the lentiviral constructs were selected with 15 μg/mL blasticidin.

### Transient transfection, protein purification, and immunoblotting

Transient transfection and cellular extract production were performed as previously described (Fu *et al*, 2010). Briefly, 2.5 x 10^6^ 293T HEK cells were transiently transfected by calcium phosphate DNA precipitation with 10-20 μg of plasmid DNA. Cells were harvested by trypsinization and washed once with PBS. The cell pellet was resuspended in 500 μL hypotonic lysis buffer (20 mM HEPES, pH 7.9; 2 mM MgCl2; 0.2 mM EGTA, 10% glycerol, 0.1 mM PMSF, 1 mM DTT), incubated on ice for 5 minutes and subjected to three freeze-thaw cycles in liquid nitrogen and 37°C. NaCl was added to extracts to a final concentration of 400 mM. After centrifugation at 14,000 x g for 15 minutes at 4°C, an equal amount of hypotonic lysis buffer with 0.2% NP-40 was added to 500 μL of soluble cellular extract.

For Strep-tag purification, whole cell extract from transiently transfected cells cell lines (1 mg of total protein) was rotated for 2 hours at 4° C in lysis buffer (20 mM HEPES at pH 7.9, 2 mM MgCl2, 0.2 mM EGTA, 10% glycerol, 1 mM DTT, 0.1 mM PMSF, 0.1% NP-40) with 200 mM NaCl. Resin was washed three times using the same buffer followed by protein analysis. Strep-tagged proteins were purified using MagSTREP “type3” XT beads, 5% suspension (IBA Lifesciences) and eluted with desthiobiotin. FLAG-tagged proteins were purified by incubating whole cell lysates from the transfected cell lines with 20 μL of DYKDDDDK-Tag Monoclonal Antibody Magnetic Microbead (Syd Labs) for three hours at 4 °C. Magnetic resin was washed three times in hypotonic lysis buffer with 200 mM NaCl.

Immunoblotting was performed as previously described (Ramos *et al*, 2019). Briefly, cell extracts and purified protein samples were boiled at 95°C for 5 minutes followed by fractionation on NuPAGE Bis-Tris polyacrylamide gels (Thermo Scientific). Separated proteins were transferred to Immobilon FL polyvinylidene difluoride (PVDF) membrane (Millipore) for immunoblotting. Membrane was blocked by Odyssey blocking buffer for 1 hour at room temperature followed by immunoblotting with the following antibodies: mouse monoclonal anti-TRMT1 aa 201-229 (G3, sc-373687, Santa Cruz Biotechnologies), rabbit polyclonal anti-TRMT1 aa 609-659 (Bethyl, A304-205A); anti-Strep-tag II (NC9261069, Thermo Fisher), anti-FLAG epitope tag (L00018; Sigma), Rabbit polyclonal anti-SARS-CoV-2 nucleoprotein N protein (40068-RP01; Sino Biological) and anti-actin (L00003; EMD Millipore). Proteins were detected using a 1:10,000 dilution of fluorescent IRDye 800CW goat anti-mouse IgG (925-32210; Thermofisher) or IRDye 680RD Goat anti-Mouse IgG Secondary Antibody (925-68070; Li-COR). Immunoblots were scanned using direct infrared fluorescence via the Odyssey system (LI-COR Biosciences).

Immunoblots were quantified and analyzed using Image Studio Version 5.2 (Li-Cor Biosciences). Rectangles of the same dimension for a given band were used to measure raw signal intensity in either the 700 or 800 nm channel. The intensity of a band in a lane was normalized to actin as a load control. For quantification of the N-terminal TRMT1 cleavage band, we measured the total signal of both the cleavage band and the non-specific band in all lanes. After normalization to actin, the total signal from the cleavage band and the non-specific band in the control lane from cells expressing GFP was subtracted from the lanes with cells expressing Nsp5 to calculate the signal arising from the cleavage band. Statistical analyses were performed using GraphPad Prism software. Where applicable, error bars represent the standard deviation. Statistical tests and the number of times each experiment was repeated are noted in each figure legend. Original files of the full raw unedited blots are available in the accompanying source data file.

### RNA analysis by primer extension

Total RNA was extracted using TRIzol LS reagent (Invitrogen). RNAs were diluted into formamide load buffer, heated to 95°C for 3 minutes, and fractionated on a 10% polyacrylamide, Tris-Borate-EDTA (TBE) gel containing 7M urea. Sybr Gold nucleic acid staining (Invitrogen) was conducted to identify the RNA pattern. For primer extension analysis, 1.5 μg of total RNA was pre-annealed with 5’-^32^P-labeled oligonucleotide and 5x hybridization buffer (250 mM Tris, pH 8.5, and 300 mM NaCl) in a total volume of 7 μl. The mixture was heated at 95°C for 3 min followed by slow cooling to 42°C. An equal amount of extension mix consisting of avian myeloblastosis virus reverse transcriptase (Promega), 5x AMV buffer and 40 μM dNTPs was added. The mixture was then incubated at 42°C for 1 hour and loaded on 15% 7M urea denaturing polyacrylamide gel. Gels were exposed on a phosphor screen (GE Healthcare) and scanned on a Bio-Rad personal molecular followed by analysis using NIH ImageJ software. Primer extension oligonucleotide sequences were previously described (Dewe *et al*., 2017). Full unedited scan images are available in the accompanying source data file.

### Subcellular localization

For localization of TRMT1 tagged with GFP, 293T human cells were seeded onto coverslips in a 6-well plate followed by transfection with the GFP plasmids noted above using Lipofectamine 3000 reagent (Thermo Fisher). For mitochondrial localization, the cells were infected with baculovirus expressing RFP targeted to mitochondria (CellLight Mitochondria-RFP, BacMam 2.0, Life Technologies). To visualize the nucleus, Hoechst dye was added to the media for 30 min before the cells were washed with PBS, fixed with 4% formaldehyde, and mounted in Aqua Poly/Mount (Polysciences Inc) followed by imaging on a Nikon A1R HD microscope.

### Assays for viral replication

The SARS-CoV-2 was a French Ile de France isolate (www.european-virus-archive.com/virus/sars-cov-2-isolate-betacovfranceidf03722020). Viral stocks were generated by amplification on VeroE6 cells. The supernatant was collected, filtered through a 0.45 µm membrane, and tittered using a TCID50 assay. For infections, the cells were previously transduced with a Lentiviral vector expressing ACE2 using the lentiviral construct RRL.sin.cPPT.SFFV/Ace2.WPRE (MT136) kindly provided by Caroline Goujon (Addgene plasmid # 145842) (Rebendenne *et al*, 2021). Seventy-two hours after transduction, accurate ACE2 expression was controlled on western blot probed with anti-ACE2 antibody (Human ACE-2 Antibody, AF933, R&D systems). ACE2-positive cells (70-80% confluence) were then infected with SARS-CoV-2 diluted to achieve the desired MOI. After 24 hours in culture, the cells were lysed with the Luna cell ready lysis module (New England Biolabs).

To measure RNA levels, the amplification reaction was run on a LightcyclerR 480 thermocycler (Roche Diagnostics) using the Luna Universal One-Step RT-qPCR kit (New England Biolabs) with the following primers:

SARS_For: 5’-ACAGGTACGTTAATAGTTAATAGCGT

SARS_Rev: 5’-ATATTGCAGCAGTACGCACACA

GAPDH_For: 5’-GCTCACCGGCATGGCCTTTCGCGT

GAPDH_Rev: 5’-TGGAGGAGTGGGTGTCGCTGTTGA.

Each qPCR was performed in triplicate, and the means and standard deviations were calculated. Relative quantification of data obtained from RT-qPCR was used to determine changes in SARS-CoV-2 Envelope (E) gene expression across multiple samples after normalization to the internal reference GAPDH gene. The raw qPCR values are provided in the source data file.

### SARS-CoV-2 infectivity measurements

Viral supernatants were collected and filtered on 0.45 µm filters. Virus was then purified by centrifugation through a 20% sucrose cushion at 25,000 rpm for 2.5 hours at 4 °C in a Sw32Ti rotor (Beckman Coulter). Virus pellets were resuspended in PBS and virus genomic RNA copy number was quantified by RTqPCR as described above. Absolute quantification of data obtained from RT-qPCR was used to determine changes in SARS-CoV-2 Envelope (E) gene expression across multiple samples by using external standards of SARS-CoV-2 (dilution 10^7^ to 10 copies) amplified in parallel within the same instrument run. The raw qPCR values are provided in the source data file. The TCID50 endpoint assay was performed on 96-well plates on VeroE6 cells using the Spearman Karber algorithm (Hierholzer & Killington, 1996).

### Source data

Unedited blot images, gels, and raw measurement values are available in the source data file associated with this manuscript and deposited onto the Dryad data repository.

## Acknowledgments

The research in this manuscript was supported by NIH T32 Training Grants GM068411 and AI1049815 to J.C.; the Montpellier University of Excellence (CoVIMOD FRS13) program to P.E. and L.B; NIH AI127370 and AI50698 to J.M.; and NSF RAPID 2033354 and NIH GR530882 to D.F. We thank Chenghong Deng, Cailyn Leo, Logan Edvalson, and Sina Ghaemmaghami for comments on this manuscript; the Mass Spectrometry Resource Lab at the University of Rochester; and the Clinical Proteomics Platform, CHU Montpellier.

## Author Contributions

K.Z., P.E., J.H.C., L.B., J.M.L., J.R., J.M., and D.F. designed the research. K.Z., P.E., J.H.C., L.B., J.M.L., J.R., and J.C. performed the research. K.Z., P.E., J.H.C., L.B., J.M.L., J.R., J.M., and D.F. analyzed the data. D.F. wrote the manuscript. K.Z., P.E., J.H.C., L.B., J.M.L., J.R., J.M., and D.F. edited the manuscript.

---

**Supplemental Figure 1.**
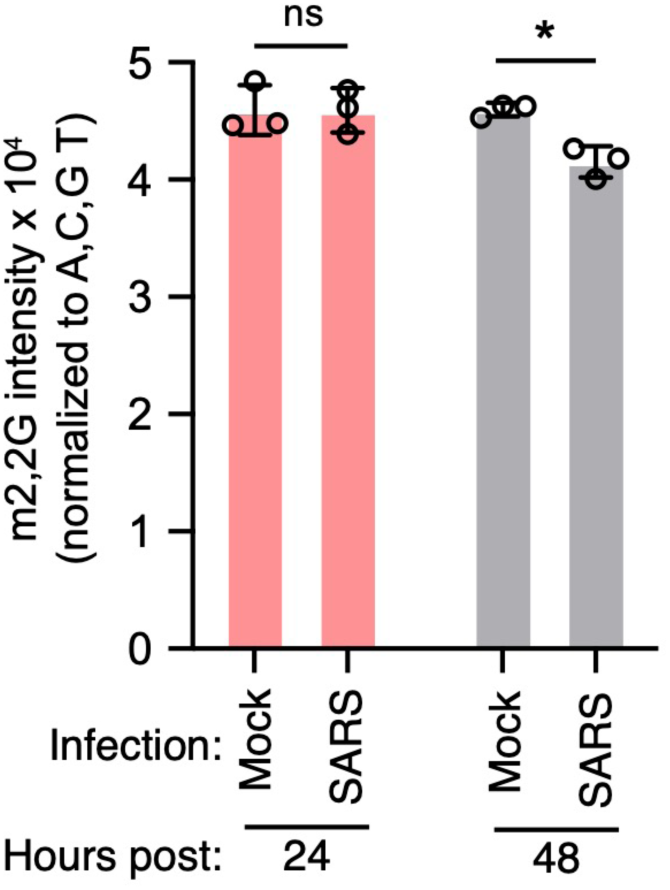
LC-MS analysis of m2,2G levels in small RNAs isolated from MRC5 cells that were either mock-infected or infected with SARS-CoV-2 at MOI of 5 for 24 or 48 hours. m2,2G levels were normalized to guanosine. Samples were measured in biological replicates. Statistical significance was determined by two-way ANOVA with multiple comparisons test. *p < 0.05; ns, non-significant.

**Supplemental Figure 2.**
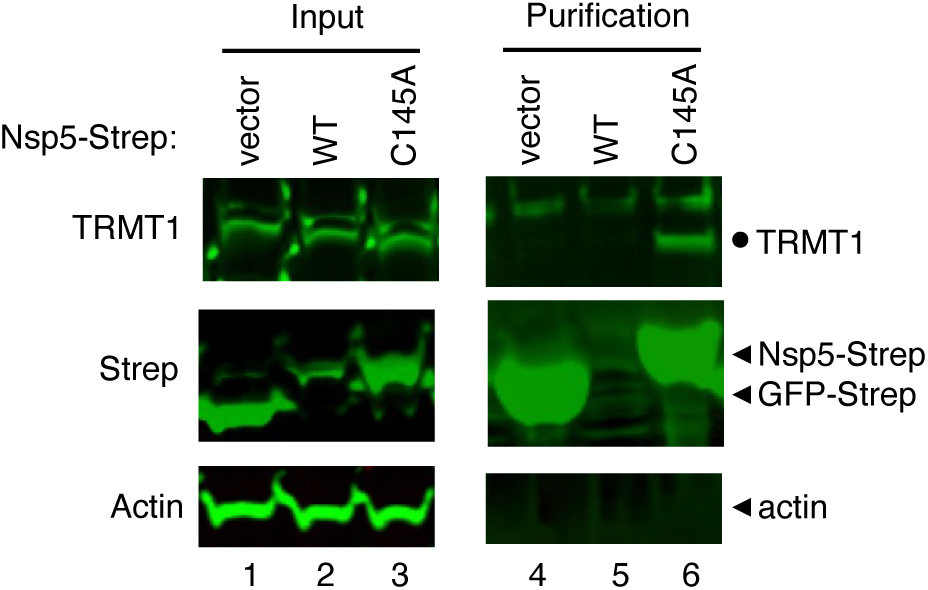
Immunoblot of input and streptactin purifications from human cells expressing empty vector, wildtype (WT) Nsp5, or Nsp5-C145A fused to the Strep tag without or with co-expression with TRMT1-FLAG. The immunoblot was probed with anti-TRMT1, Strep, or actin antibodies. Circle represents endogenous TRMT1.

**Supplemental Figure 3.**
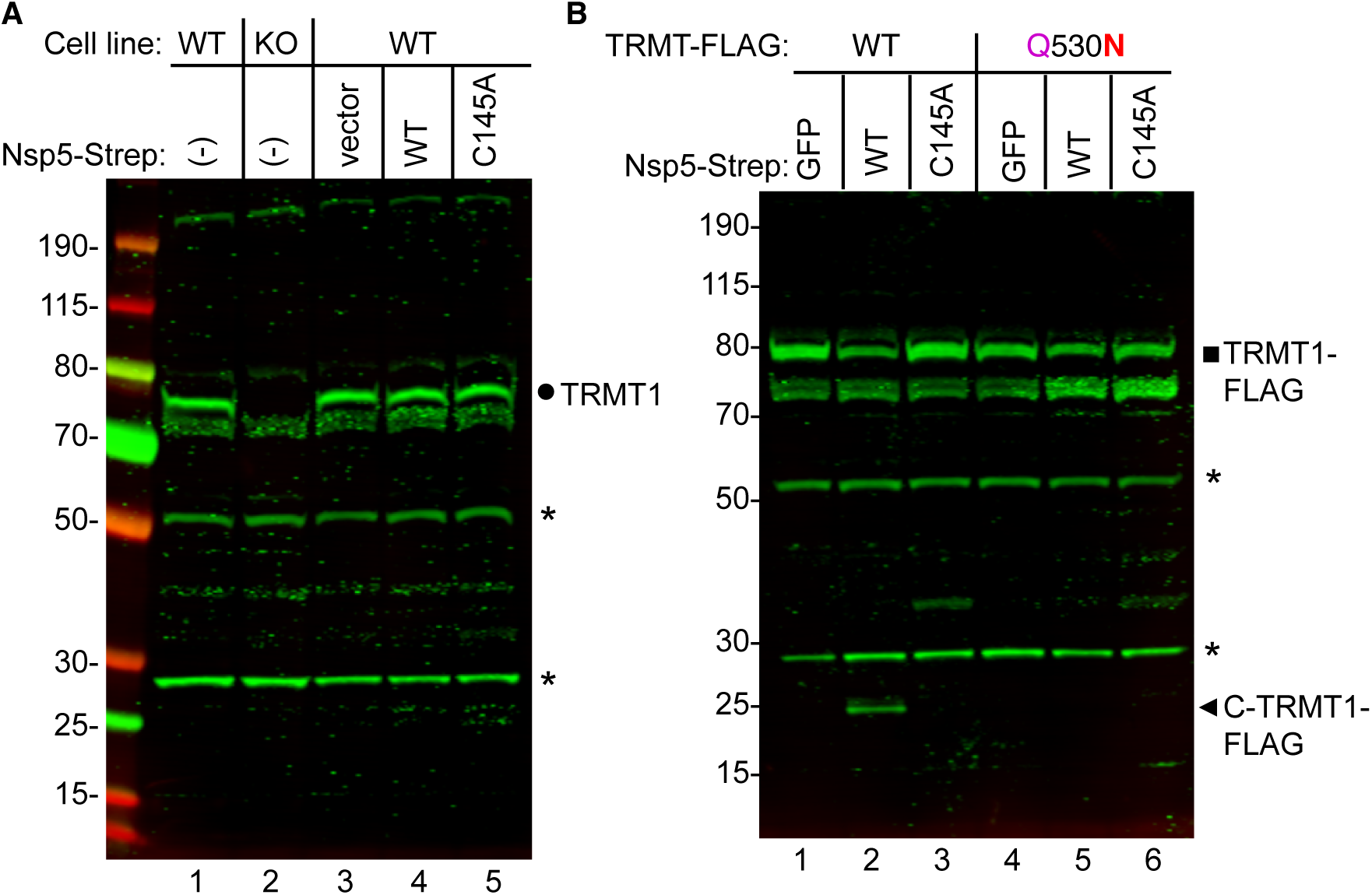
Detection of the C-terminal TRMT1 fragment produced by Nsp5-dependent cleavage in human cells. (A) Immunoblot of lysates from the indicated wildtype (WT) or TRMT1-KO human 293T cells that were untransfected (-) (lanes 1 and 2) or transfected with empty vector, wildtype (WT) Nsp5-Strep, or Nsp5-C145A-Strep expression plasmids (lanes 3 through 5). The blot was probed with an antibody detecting residues 609-659 of TRMT1. Circle represents endogenous TRMT1. (B) Immunoblot of lysates from human cells that were transfected with TRMT1-FLAG or TRMT1-FLAG Q530N expression plasmids along with empty vector, wildtype (WT) Nsp5-Strep, or Nsp5-C145A-Strep expression plasmids. The blot was probed with an antibody detecting residues 609-659 of TRMT1. Circle represents endogenous TRMT1, square represents TRMT1-FLAG, and arrowhead indicates the C-terminal TRMT1 cleavage product. Asterisks denote bands that are still detectable in the TRMT1-KO cell line. The band at ∼35 kDa in lanes 3 and 6 of Supplemental Figure 3B represents non-specific detection of the Nsp5-C145A variant that exhibits extremely high levels of expression since it cannot self-cleave.

**Supplemental Figure 4.**
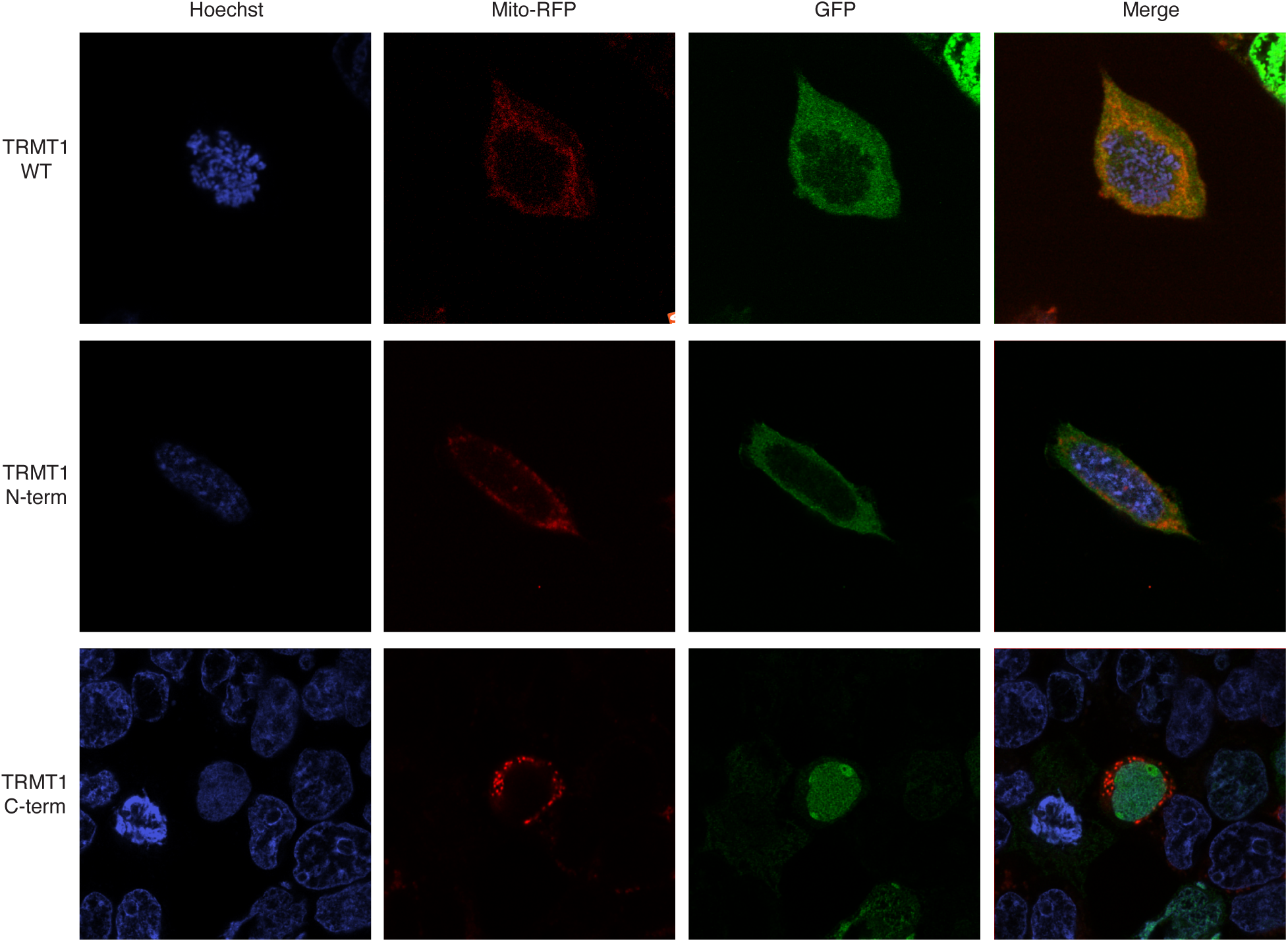
Confocal microscopy images of 293T cells transiently transfected with constructs expressing TRMT1 or TRMT1 fragments fused with green fluorescent protein (GFP). Mitochondria were identified using mitochondrion-targeted red fluorescent protein (Mito-RFP) and nuclear DNA was stained with Hoechst. Overlap of red mitochondria and green GFP signal is displayed in the Merge panels.

**Supplemental Figure 5.**
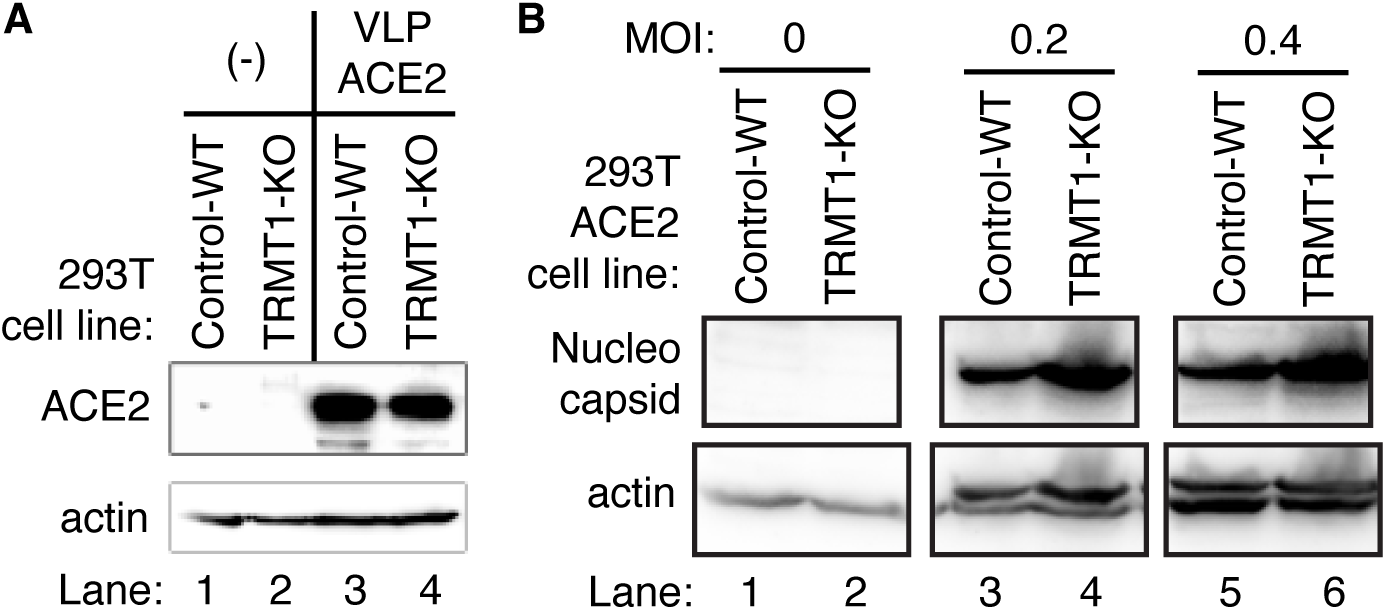
Human 293T cell lines expressing ACE2 can be infected by SARS-CoV-2. (A) Immunoblot analysis of lysates prepared from the indicated 293T cell lines expressing empty vector or ACE2. The immunoblot was probed with anti-ACE2 and actin. (B) Immunoblot analysis of lysates prepared from 293T-ACE2 cell lines that were mock-infected (MOI 0) or infected with SARS-CoV-2 at the indicated MOI. The blot was probed against the SARS-CoV-2 nucleocapsid (N) and actin.

**Supplemental Figure 6.**
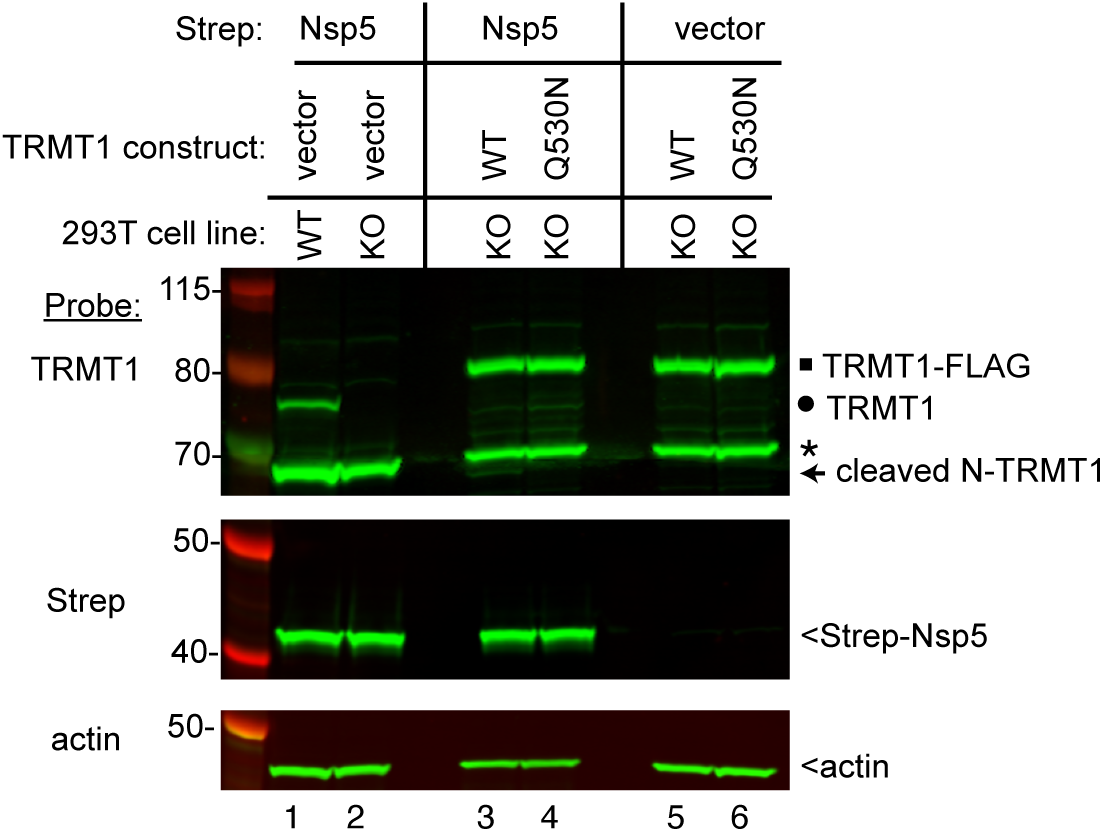
Expression of Nsp5 leads to cleavage of TRMT1-WT re-expressed in TRMT1-KO cells, but not TRMT1-Q530N. Immunoblot of lysates from human cells integrated with empty lentiviral vector or lentiviral expression vectors for wildtype (WT) TRMT1 or TRMT1-Q530N. The cell lines were transfected with either vector or a construct expressing Nsp5-Strep. The immunoblot was probed with anti-TRMT1, Strep, and actin antibodies. Square represents TRMT1-FLAG, circle represent endogenous TRMT1, * denotes a non-specific band, and arrow represents N-terminal TRMT1 cleavage product.

**Supplemental Figure 7.**
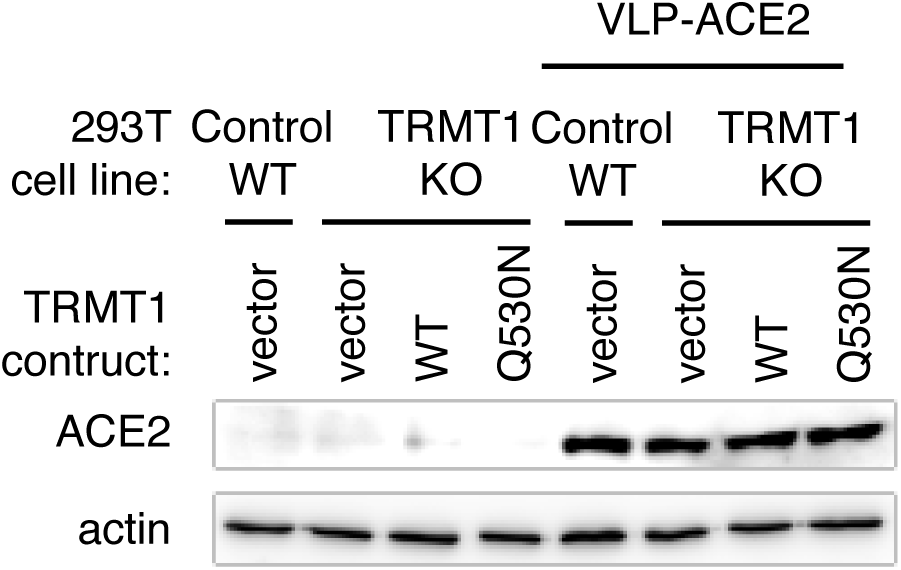
Immunoblot analysis of lysates prepared from the indicated 293T cell lines expressing empty vector or ACE2. The immunoblot was probed with anti-ACE2 and actin.

## References

Baumstark T, Ahlquist P (2001) The brome mosaic virus RNA3 intergenic replication enhancer folds to mimic a tRNA TpsiC-stem loop and is modified in vivo. RNA 7: 1652–1670

Becker HF, Motorin Y, Florentz C, Giege R, Grosjean H (1998) Pseudouridine and ribothymidine formation in the tRNA-like domain of turnip yellow mosaic virus RNA. Nucleic Acids Res 26: 3991–3997

Blaesius K, Abbasi AA, Tahir TH, Tietze A, Picker-Minh S, Ali G, Farooq S, Hu H, Latif Z, Khan MN, Kaindl A (2018) Mutations in the tRNA methyltransferase 1 gene TRMT1 cause congenital microcephaly, isolated inferior vermian hypoplasia and cystic leukomalacia in addition to intellectual disability. Am J Med Genet A 176: 2517–2521

Bojkova D, Klann K, Koch B, Widera M, Krause D, Ciesek S, Cinatl J, Munch C (2020) Proteomics of SARS-CoV-2-infected host cells reveals therapy targets. Nature 583: 469–472

Burgess HM, Depledge DP, Thompson L, Srinivas KP, Grande RC, Vink EI, Abebe JS, Blackaby WP, Hendrick A, Albertella MR et al (2021) Targeting the m(6)A RNA modification pathway blocks SARS-CoV-2 and HCoV-OC43 replication. Genes Dev 35: 1005–1019

Cai WM, Chionh YH, Hia F, Gu C, Kellner S, McBee ME, Ng CS, Pang YL, Prestwich EG, Lim KS et al (2015) A Platform for Discovery and Quantification of Modified Ribonucleosides in RNA: Application to Stress-Induced Reprogramming of tRNA Modifications. Methods Enzymol 560: 29–71

Chan C, Pham P, Dedon PC, Begley TJ (2018) Lifestyle modifications: coordinating the tRNA epitranscriptome with codon bias to adapt translation during stress responses. Genome Biol 19: 228

Chen CC, Yu X, Kuo CJ, Min J, Chen S, Ma L, Liu K, Guo RT (2021) Overview of antiviral drug candidates targeting coronaviral 3C-like main proteases. FEBS J 288: 5089–5121

D’Oliviera A, Dai X, Mottaghinia S, Geissler EP, Etienne L, Zhang Y, Mugridge JS (2023) Recognition and Cleavage of Human tRNA Methyltransferase TRMT1 by the SARS-CoV-2 Main Protease. bioRxiv: 2023.2002.2020.529306

D′Oliviera A, Dai X, Mottaghinia S, Geissler EP, Etienne L, Zhang Y, Mugridge JS, 2023. Recognition and Cleavage of Human tRNA Methyltransferase TRMT1 by the SARS-CoV-2 Main Protease. eLife Sciences Publications, Ltd.

Dewe JM, Fuller BL, Lentini JM, Kellner SM, Fu D (2017) TRMT1-Catalyzed tRNA Modifications Are Required for Redox Homeostasis To Ensure Proper Cellular Proliferation and Oxidative Stress Survival. Mol Cell Biol 37

Di Giorgio S, Martignano F, Torcia MG, Mattiuz G, Conticello SG (2020) Evidence for host-dependent RNA editing in the transcriptome of SARS-CoV-2. Sci Adv 6: eabb5813

Dremel SE, Jimenez AR, Tucker JM (2023) “Transfer” of power: The intersection of DNA virus infection and tRNA biology. Semin Cell Dev Biol

Dremel SE, Sivrich FL, Tucker JM, Glaunsinger BA, DeLuca NA (2022) Manipulation of RNA polymerase III by Herpes Simplex Virus-1. Nat Commun 13: 623

Finkel Y, Gluck A, Nachshon A, Winkler R, Fisher T, Rozman B, Mizrahi O, Lubelsky Y, Zuckerman B, Slobodin B et al (2021) SARS-CoV-2 uses a multipronged strategy to impede host protein synthesis. Nature 594: 240–245

Fu D, Brophy JA, Chan CT, Atmore KA, Begley U, Paules RS, Dedon PC, Begley TJ, Samson LD (2010) Human AlkB homolog ABH8 Is a tRNA methyltransferase required for wobble uridine modification and DNA damage survival. Mol Cell Biol 30: 2449–2459

Gordon DE, Hiatt J, Bouhaddou M, Rezelj VV, Ulferts S, Braberg H, Jureka AS, Obernier K, Guo JZ, Batra J et al (2020a) Comparative host-coronavirus protein interaction networks reveal pan-viral disease mechanisms. Science 370

Gordon DE, Jang GM, Bouhaddou M, Xu J, Obernier K, White KM, O’Meara MJ, Rezelj VV, Guo JZ, Swaney DL et al (2020b) A SARS-CoV-2 protein interaction map reveals targets for drug repurposing. Nature 583: 459–468

Grum-Tokars V, Ratia K, Begaye A, Baker SC, Mesecar AD (2008) Evaluating the 3C-like protease activity of SARS-Coronavirus: recommendations for standardized assays for drug discovery. Virus Res 133: 63–73

Hartenian E, Nandakumar D, Lari A, Ly M, Tucker JM, Glaunsinger BA (2020) The molecular virology of coronaviruses. J Biol Chem 295: 12910–12934

Heilmann E, Costacurta F, Geley S, Mogadashi SA, Volland A, Rupp B, Harris RS, von Laer D (2022) A VSV-based assay quantifies coronavirus Mpro/3CLpro/Nsp5 main protease activity and chemical inhibition. Commun Biol 5: 391

Hernandez-Alias X, Benisty H, Schaefer MH, Serrano L (2021) Translational adaptation of human viruses to the tissues they infect. Cell Rep 34: 108872

Hierholzer J, Killington R (1996) Virus isolation and quantitation. In: Virology methods manual, pp. 25-46. Elsevier:

Jin D, Musier-Forsyth K (2019) Role of host tRNAs and aminoacyl-tRNA synthetases in retroviral replication. J Biol Chem 294: 5352–5364

Jin Y, Ouyang M, Yu T, Zhuang J, Wang W, Liu X, Duan F, Guo D, Peng X, Pan JA (2022) Genome-Wide Analysis of the Indispensable Role of Non-structural Proteins in the Replication of SARS-CoV-2. Front Microbiol 13: 907422

Jonkhout N, Cruciani S, Santos Vieira HG, Tran J, Liu H, Liu G, Pickford R, Kaczorowski D, Franco GR, Vauti F et al (2021) Subcellular relocalization and nuclear redistribution of the RNA methyltransferases TRMT1 and TRMT1L upon neuronal activation. RNA Biol 18: 1905–1919

Jumper J, Evans R, Pritzel A, Green T, Figurnov M, Ronneberger O, Tunyasuvunakool K, Bates R, Zidek A, Potapenko A et al (2021) Highly accurate protein structure prediction with AlphaFold. Nature 596: 583–589

Jungfleisch J, Bottcher R, Tallo-Parra M, Perez-Vilaro G, Merits A, Novoa EM, Diez J (2022) CHIKV infection reprograms codon optimality to favor viral RNA translation by altering the tRNA epitranscriptome. Nat Commun 13: 4725

Kamitani W, Huang C, Narayanan K, Lokugamage KG, Makino S (2009) A two-pronged strategy to suppress host protein synthesis by SARS coronavirus Nsp1 protein. Nat Struct Mol Biol 16: 1134–1140

Katanski CD, Alshammary H, Watkins CP, Huang S, Gonzales-Reiche A, Sordillo EM, van Bakel H, Mount Sinai PSPsg, Lolans K, Simon V, Pan T (2022) tRNA abundance, modification and fragmentation in nasopharyngeal swabs as biomarkers for COVID-19 severity. Front Cell Dev Biol 10: 999351

Kim D, Lee JY, Yang JS, Kim JW, Kim VN, Chang H (2020) The Architecture of SARS-CoV-2 Transcriptome. Cell 181: 914–921 e910

Lamers MM, Haagmans BL (2022) SARS-CoV-2 pathogenesis. Nat Rev Microbiol 20: 270–284

Lapointe CP, Grosely R, Johnson AG, Wang J, Fernandez IS, Puglisi JD (2021) Dynamic competition between SARS-CoV-2 NSP1 and mRNA on the human ribosome inhibits translation initiation. Proc Natl Acad Sci U S A 118

Lee J, Kenward C, Worrall LJ, Vuckovic M, Gentile F, Ton AT, Ng M, Cherkasov A, Strynadka NCJ, Paetzel M (2022) X-ray crystallographic characterization of the SARS-CoV-2 main protease polyprotein cleavage sites essential for viral processing and maturation. Nat Commun 13: 5196

Lee J, Worrall LJ, Vuckovic M, Rosell FI, Gentile F, Ton AT, Caveney NA, Ban F, Cherkasov A, Paetzel M, Strynadka NCJ (2020) Crystallographic structure of wild-type SARS-CoV-2 main protease acyl-enzyme intermediate with physiological C-terminal autoprocessing site. Nat Commun 11: 5877

Lesiewicz J, Dudock B (1978) In vitro methylation of tobacco mosaic virus RNA with ribothymidine-forming tRNA methyltransferase. Characterization and specificity of the reaction. Biochim Biophys Acta 520: 411–418

Li C, Yao Y, Long D, Lin X (2021a) KDELC1 and TRMT1 Serve as Prognosis-Related SARS-CoV-2 Proteins Binding Human mRNAs and Promising Biomarkers in Clear Cell Renal Cell Carcinoma. Int J Gen Med 14: 2475–2490

Li N, Hui H, Bray B, Gonzalez GM, Zeller M, Anderson KG, Knight R, Smith D, Wang Y, Carlin AF, Rana TM (2021b) METTL3 regulates viral m6A RNA modification and host cell innate immune responses during SARS-CoV-2 infection. Cell Rep 35: 109091

Liu Y, Qin C, Rao Y, Ngo C, Feng JJ, Zhao J, Zhang S, Wang TY, Carriere J, Savas AC et al (2021) SARS-CoV-2 Nsp5 Demonstrates Two Distinct Mechanisms Targeting RIG-I and MAVS To Evade the Innate Immune Response. mBio 12: e0233521

Lu JL, Zhou XL (2023) SARS-CoV-2 main protease Nsp5 cleaves and inactivates human tRNA methyltransferase TRMT1. J Mol Cell Biol

Merad M, Blish CA, Sallusto F, Iwasaki A (2022) The immunology and immunopathology of COVID-19. Science 375: 1122–1127

Meyer B, Chiaravalli J, Gellenoncourt S, Brownridge P, Bryne DP, Daly LA, Grauslys A, Walter M, Agou F, Chakrabarti LA et al (2021) Characterising proteolysis during SARS-CoV-2 infection identifies viral cleavage sites and cellular targets with therapeutic potential. Nat Commun 12: 5553

Meyers JM, Ramanathan M, Shanderson RL, Beck A, Donohue L, Ferguson I, Guo MG, Rao DS, Miao W, Reynolds D et al (2021) The proximal proteome of 17 SARS-CoV-2 proteins links to disrupted antiviral signaling and host translation. PLoS Pathog 17: e1009412

Minkoff JM, Tenoever B (2023) Innate immune evasion strategies of SARS-CoV-2. Nat Rev Microbiol 21: 178–194

Moustaqil M, Ollivier E, Chiu HP, Van Tol S, Rudolffi-Soto P, Stevens C, Bhumkar A, Hunter DJB, Freiberg AN, Jacques D et al (2021) SARS-CoV-2 proteases PLpro and 3CLpro cleave IRF3 and critical modulators of inflammatory pathways (NLRP12 and TAB1): implications for disease presentation across species. Emerg Microbes Infect 10: 178–195

Muramatsu T, Kim YT, Nishii W, Terada T, Shirouzu M, Yokoyama S (2013) Autoprocessing mechanism of severe acute respiratory syndrome coronavirus 3C-like protease (SARS-CoV 3CLpro) from its polyproteins. FEBS J 280: 2002–2013

Najmabadi H, Hu H, Garshasbi M, Zemojtel T, Abedini SS, Chen W, Hosseini M, Behjati F, Haas S, Jamali P et al (2011) Deep sequencing reveals 50 novel genes for recessive cognitive disorders. Nature 478: 57–63

Narayanan A, Toner SA, Jose J (2022) Structure-based inhibitor design and repurposing clinical drugs to target SARS-CoV-2 proteases. Biochem Soc Trans 50: 151–165

Netzer N, Goodenbour JM, David A, Dittmar KA, Jones RB, Schneider JR, Boone D, Eves EM, Rosner MR, Gibbs JS et al (2009) Innate immune and chemically triggered oxidative stress modifies translational fidelity. Nature 462: 522–526

Nie X, Qian L, Sun R, Huang B, Dong X, Xiao Q, Zhang Q, Lu T, Yue L, Chen S et al (2021) Multi-organ proteomic landscape of COVID-19 autopsies. Cell 184: 775–791 e714

Nunes A, Ribeiro DR, Marques M, Santos MAS, Ribeiro D, Soares AR (2020) Emerging Roles of tRNAs in RNA Virus Infections. Trends Biochem Sci 45: 794–805

Owen DR, Allerton CMN, Anderson AS, Aschenbrenner L, Avery M, Berritt S, Boras B, Cardin RD, Carlo A, Coffman KJ et al (2021) An oral SARS-CoV-2 M(pro) inhibitor clinical candidate for the treatment of COVID-19. Science 374: 1586–1593

Pallan PS, Kreutz C, Bosio S, Micura R, Egli M (2008) Effects of N2,N2-dimethylguanosine on RNA structure and stability: crystal structure of an RNA duplex with tandem m2 2G:A pairs. RNA 14: 2125–2135

Pavon-Eternod M, David A, Dittmar K, Berglund P, Pan T, Bennink JR, Yewdell JW (2013) Vaccinia and influenza A viruses select rather than adjust tRNAs to optimize translation. Nucleic Acids Res 41: 1914–1921

Pena N, Zhang W, Watkins C, Halucha M, Alshammary H, Hernandez MM, Liu WC, Albrecht RA, Garcia-Sastre A, Simon V et al (2022) Profiling Selective Packaging of Host RNA and Viral RNA Modification in SARS-CoV-2 Viral Preparations. Front Cell Dev Biol 10: 768356

Peng X, Luo Y, Li H, Guo X, Chen H, Ji X, Liang H (2022) RNA editing increases the nucleotide diversity of SARS-CoV-2 in human host cells. PLoS Genet 18: e1010130

Perez M, Nance KD, Bak DW, Thalalla Gamage S, Najera SS, Conte AN, Linehan WM, Weerapana E, Meier JL (2022) Conditional Covalent Lethality Driven by Oncometabolite Accumulation. ACS Chem Biol 17: 2789–2800

Pettersen EF, Goddard TD, Huang CC, Couch GS, Greenblatt DM, Meng EC, Ferrin TE (2004) UCSF Chimera--a visualization system for exploratory research and analysis. J Comput Chem 25: 1605–1612

Ramos J, Han L, Li Y, Hagelskamp F, Kellner SM, Alkuraya FS, Phizicky EM, Fu D (2019) Formation of tRNA Wobble Inosine in Humans Is Disrupted by a Millennia-Old Mutation Causing Intellectual Disability. Molecular and cellular biology 39

Raymonda MH, Ciesla JH, Monaghan M, Leach J, Asantewaa G, Smorodintsev-Schiller LA, Lutz MMt, Schafer XL, Takimoto T, Dewhurst S et al (2022) Pharmacologic profiling reveals lapatinib as a novel antiviral against SARS-CoV-2 in vitro. Virology 566: 60–68

Rebendenne A, Valadao ALC, Tauziet M, Maarifi G, Bonaventure B, McKellar J, Planes R, Nisole S, Arnaud-Arnould M, Moncorge O, Goujon C (2021) SARS-CoV-2 triggers an MDA-5-dependent interferon response which is unable to control replication in lung epithelial cells. J Virol 95

Resnick SJ, Iketani S, Hong SJ, Zask A, Liu H, Kim S, Melore S, Lin FY, Nair MS, Huang Y et al (2021) Inhibitors of Coronavirus 3CL Proteases Protect Cells from Protease-Mediated Cytotoxicity. J Virol 95: e0237420

Samavarchi-Tehrani P, Abdouni H, Knight JDR, Astori A, Samson R, Lin Z-Y, Kim D-K, Knapp JJ, St-Germain J, Go CD et al (2020) A SARS-CoV-2 – host proximity interactome. bioRxiv: 2020.2009.2003.282103

Schmidt TG, Batz L, Bonet L, Carl U, Holzapfel G, Kiem K, Matulewicz K, Niermeier D, Schuchardt I, Stanar K (2013) Development of the Twin-Strep-tag(R) and its application for purification of recombinant proteins from cell culture supernatants. Protein Expr Purif 92: 54–61

Schubert K, Karousis ED, Jomaa A, Scaiola A, Echeverria B, Gurzeler LA, Leibundgut M, Thiel V, Muhlemann O, Ban N (2020) SARS-CoV-2 Nsp1 binds the ribosomal mRNA channel to inhibit translation. Nat Struct Mol Biol 27: 959–966

Steinberg S, Cedergren R (1995) A correlation between N2-dimethylguanosine presence and alternate tRNA conformers. RNA 1: 886–891

Suryawanshi RK, Koganti R, Agelidis A, Patil CD, Shukla D (2021) Dysregulation of Cell Signaling by SARS-CoV-2. Trends Microbiol 29: 224–237

Thoms M, Buschauer R, Ameismeier M, Koepke L, Denk T, Hirschenberger M, Kratzat H, Hayn M, Mackens-Kiani T, Cheng J et al (2020) Structural basis for translational shutdown and immune evasion by the Nsp1 protein of SARS-CoV-2. Science 369: 1249–1255

Tidu A, Janvier A, Schaeffer L, Sosnowski P, Kuhn L, Hammann P, Westhof E, Eriani G, Martin F (2020) The viral protein NSP1 acts as a ribosome gatekeeper for shutting down host translation and fostering SARS-CoV-2 translation. RNA 27: 253–264

Tucker JM, Schaller AM, Willis I, Glaunsinger BA (2020) Alteration of the Premature tRNA Landscape by Gammaherpesvirus Infection. mBio 11

Uemura K, Nobori H, Sato A, Sanaki T, Toba S, Sasaki M, Murai A, Saito-Tarashima N, Minakawa N, Orba Y et al (2021a) 5-Hydroxymethyltubercidin exhibits potent antiviral activity against flaviviruses and coronaviruses, including SARS-CoV-2. iScience 24: 103120

Uemura K, Sasaki M, Sanaki T, Toba S, Takahashi Y, Orba Y, Hall WW, Maenaka K, Sawa H, Sato A (2021b) MRC5 cells engineered to express ACE2 serve as a model system for the discovery of antivirals targeting SARS-CoV-2. Sci Rep 11: 5376

V’Kovski P, Kratzel A, Steiner S, Stalder H, Thiel V (2021) Coronavirus biology and replication: implications for SARS-CoV-2. Nat Rev Microbiol 19: 155–170

van Weringh A, Ragonnet-Cronin M, Pranckeviciene E, Pavon-Eternod M, Kleiman L, Xia X (2011) HIV-1 modulates the tRNA pool to improve translation efficiency. Mol Biol Evol 28: 1827–1834

Varadi M, Anyango S, Deshpande M, Nair S, Natassia C, Yordanova G, Yuan D, Stroe O, Wood G, Laydon A et al (2022) AlphaFold Protein Structure Database: massively expanding the structural coverage of protein-sequence space with high-accuracy models. Nucleic Acids Res 50: D439–D444

Wenzel J, Lampe J, Muller-Fielitz H, Schuster R, Zille M, Muller K, Krohn M, Korbelin J, Zhang L, Ozorhan U et al (2021) The SARS-CoV-2 main protease M(pro) causes microvascular brain pathology by cleaving NEMO in brain endothelial cells. Nat Neurosci 24: 1522–1533

Xia Y, Li K, Li J, Wang T, Gu L, Xun L (2019) T5 exonuclease-dependent assembly offers a low-cost method for efficient cloning and site-directed mutagenesis. Nucleic Acids Res 47: e15

Yan S, Wu G (2021) Spatial and temporal roles of SARS-CoV PL(pro) −A snapshot. FASEB J 35: e21197

Yang JY, Fang W, Miranda-Sanchez F, Brown JM, Kauffman KM, Acevero CM, Bartel DP, Polz MF, Kelly L (2021) Degradation of host translational machinery drives tRNA acquisition in viruses. Cell Syst 12: 771–779 e775

Zhang K, Lentini JM, Prevost CT, Hashem MO, Alkuraya FS, Fu D (2020) An intellectual disability-associated missense variant in TRMT1 impairs tRNA modification and reconstitution of enzymatic activity. Hum Mutat 41: 600–607

Zhang LS, Xiong QP, Pena Perez S, Liu C, Wei J, Le C, Zhang L, Harada BT, Dai Q, Feng X et al (2021a) ALKBH7-mediated demethylation regulates mitochondrial polycistronic RNA processing. Nat Cell Biol 23: 684–691

Zhang S, Wang J, Cheng G (2021b) Protease cleavage of RNF20 facilitates coronavirus replication via stabilization of SREBP1. Proc Natl Acad Sci U S A 118

Zhang X, Hao H, Ma L, Zhang Y, Hu X, Chen Z, Liu D, Yuan J, Hu Z, Guan W (2021c) Methyltransferase-like 3 Modulates Severe Acute Respiratory Syndrome Coronavirus-2 RNA N6-Methyladenosine Modification and Replication. mBio 12: e0106721

